# Genomic and Behavioral Signatures of Selection for Ethanol Preference from the Heterogeneous Stock Collaborative Cross Mice – The Central Nucleus of the Amygdala

**DOI:** 10.1101/2025.04.15.649004

**Authors:** Justin Q. Anderson, Priscila Darakjian, Robert Hitzemann, Rita Cervera-Juanes, Kip D. Zimmerman, Cheryl Reed, Denesa Lockwood, Angela R. Ozburn, Tamara J. Phillips

**Affiliations:** Portland Alcohol Research Center, Department of Behavioral Neuroscience, Oregon Health & Science University, Portland, OR 97239, USA; Wake Forest University School of Medicine, Department of Translational Neuroscience, Winston-Salem, NC 27157, USA; Wake Forest University School of Medicine, Center for Precision Medicine, Winston-Salem, NC 27157, USA; Wake Forest University School of Medicine, Department of Internal Medicine, Winston-Salem, NC 27157, USA; Wake Forest University School of Medicine, Department of Biostatistics and Data Science, Winston-Salem, NC 27157, USA; Department of Neurology, Oregon Health & Science University, Portland, OR 97239, USA; VA Portland Health Care System, Portland, OR, 97239, USA

## Abstract

Alcohol use disorder (AUD) is a complex disease with heritability of ∼0.5, indicating genetic and non-genetic factors contribute to risk. Identifying gene expression networks contributing to risk using post-mortem human brain tissue has the limitation of conflating risk for AUD with consequences of alcohol use. We leveraged mice selectively bred for differential ethanol preference from a highly genetically diverse population to overcome this limitation. Ethanol intake was highly correlated with preference, high-preferring (HP) mice consumed more sweet-but not bitter-tasting solutions compared to low-preferring (LP) mice, and the lines did not differ in rate of ethanol elimination. Adult, ethanol-naïve HP and LP mice contributed tissue from the central nucleus of the amygdala (CeA), a region critical to ethanol preference and intake. Single-nuclei and bulk RNA sequencing data were used to identify cell types and transcriptome changes related to selective breeding for differential risk for ethanol preference. Single nuclei analysis identified populations of inhibitory (∼48% of cells) and excitatory (∼23%) neurons, and non-neuronal (∼29%) cells, but no differences in cell-type composition or gene expression were identified between the lines. Bulk CeA analysis identified differences between the lines for: (1) gene expression (2996 genes), (2) expression variability (426 genes), and (3) wiring (407 significant gene-gene correlations). Overall, lower variance was found in the HP line. Reduced gene-gene correlation, also found in HP mice, suggested that selection for high preference induced changes in transcriptional regulation resulting in reduced connectivity, specific to gene networks enriched in markers for inhibitory neurons expressing *Isl1* and *Tac1*.

## 1 Introduction

Human and murine populations have contributed considerable information about brain circuitry and gene networks relevant to alcohol drinking and alcohol use disorder (AUD). Dysregulation of the transcriptome preceding exposure to alcohol may contribute to risk for AUD. Using postmortem human brain tissue, Farris et al. [1] described a multi-omics approach that included RNA-Seq assessment of co-expression networks, functional annotation of the data into ontological categories, examination of gene modules for overrepresentation of genetic variants within the database of genotypes and phenotypes (dbGaP [2]), and results from a meta-analysis of alcohol self-administration in mice [1, 3]. The analysis implicated genetic factors involved in synaptic regulation. A more recent report by Rao et al. [4] reflects advances in the field allowing for identification of functionally relevant gene variants. Differences identified from post-mortem human tissue could be related to risk for AUD or to the consequences of alcohol use. Murine selective breeding provides a model of high vs. low genetic risk for alcohol (ethanol) intake in which the risk-related transcriptome can be analyzed in the absence of ethanol exposure [5–8].

Colville et al. [7] selectively bred mice for high preference (HP) or low preference (LP) for ethanol using the standard 2-bottle choice model, first described in mice by McClearn and Rodgers [9]. The relevance of this model to our understanding of AUD has been recently reviewed [10]. The concern is frequently raised that with the preference model, the animals either do not, or only briefly, exceed intoxicating blood ethanol concentrations (BECs) of 80 mg/dl. However, there is considerable evidence that the preference model can lead to substantial and persistent BECs [11–13]. There are also strategies used for selective breeding to consider. Short-term bi-directional selective breeding focuses on finding significant differences between the selected lines (e.g. 2 or 3 standard deviations), while minimizing the random fixation of genes unrelated to the selection phenotype by breeding for few generations. Short-term selective breeding can utilize a larger number of selection families to reduce inbreeding. Examples are Colville et al. [7] and Kozell et al. [14]. Long-term bi-directional selective breeding focuses on finding the most robust differences possible between the lines by breeding for many generations, essentially maximizing the between-line, selection trait-relevant genetic diversity. However, the disadvantage is that the threshold for detecting significant effects is significantly elevated given the amount of inbreeding that occurs across generations.

Colville et al. [7] used heterogeneous stock – collaborative cross (HS-CC) mice as the founder population for ethanol preference selection. The HS-CC population was derived by intercrossing 8 inbred mouse strains and encompasses ∼ 90% of the genetic diversity available in *Mus Musculus* [15]. Among the 8 strains are 2 with high ethanol preference and intake (C57BL/6 [B6] and PWK) [16]. We have reported associations of neuroimmune mechanisms with level of preference in comparisons of the high and low preference strains [17]. Colville et al. [7, 8] used RNA-Seq to examine how selective breeding for ethanol preference affected the transcriptome in three brain regions: the nucleus accumbens shell, prelimbic cortex and central nucleus of the amygdala (CeA). Although many of the transcriptional changes were region specific, some were not. A key finding was that selection affected the expression of synaptic tethering proteins involved in the regulation of glutamate receptor plasticity and in the expression of cell adhesion molecules, especially the cadherins and protocadherins.

In the present study, we build upon this foundational work to provide a robust, expanded transcriptomic analysis of replicate HP and LP lines, deepening analysis of the CeA, and integrating cell-type–specific insights. The second replicate set of HP and LP lines were bred from the HS-CC, using the same design for selective breeding as that used by Colville et al. [7].

Additional behavioral and physiological data for the selected lines were also obtained, including preference and intake for sweet and bitter tasting fluids, locomotor and hypothermic responses to ethanol and rate of ethanol elimination. The experimental design for transcriptome analysis also mirrored Colville et al. [7, 8], with some modifications: (1) The sample size for the RNA-Seq analyses was increased more than 100%. This modification was particularly important for separate analysis of male and female data. Evidence that males and females respond differently to the effects of ethanol has been reviewed elsewhere [19]. (2) Although samples were collected from multiple brain regions, we chose to initially analyze data and report results for the CeA, consistent with the initial report by Colville et al. [7]. (3) We expanded transcriptomic analysis to include single nucleus RNA-Seq (snRNA-Seq) for the CeA in both HP and LP lines, enabling us to assess cell-type composition and to assign gene co-expression modules to specific cell populations using weighted gene co-expression network analysis [WGCNA; [20, 21]). While we do not directly compare the first and second HP and LP lines here, the use of replicate lines strengthens the broader framework for identifying reproducible versus line-specific genetic signatures and contributes to evaluating the robustness of trait-associated transcriptomic profiles.

## 2 Results

### 2.1 Selective Breeding

#### 2.1.1 Ethanol Preference

Selective breeding was effective at producing lines with divergent ethanol preference, consistent with our prior work [7]. Results for generation S4, the final selection generation, are detailed here (Figure 1A). Mice from the S4 generation were used to generate those tested for other traits and for the transcriptome analyses presented here. Preference ratio data were examined by ANOVA, with ethanol concentration as a repeated measure (5 and 10% v/v ethanol), and selected line and sex as independent factors. Data are not shown by sex, because there was no significant main effect of sex, nor interaction of sex with line or concentration. There was a significant line by ethanol concentration interaction (F[1,167]=3.9, p<0.05). Simple main effects analysis identified a significant line difference in preference for both 5 and 10% ethanol (ps<0.001), with greater preference in the HP line. There was no impact of ethanol concentration on intake within the LP mice; however, the HP mice had a greater mean preference ratio for 5% ethanol, compared to 10% ethanol (p<0.001). The mean ± SEM ethanol preference values for the 5 and 10% ethanol concentrations are also shown in Figure 1A for the subset of S4 mice chosen to serve as S5 breeders. A subset of the highest and lowest preference S4 mice was also used for the single nuclei analyses.

**Figure 1.**
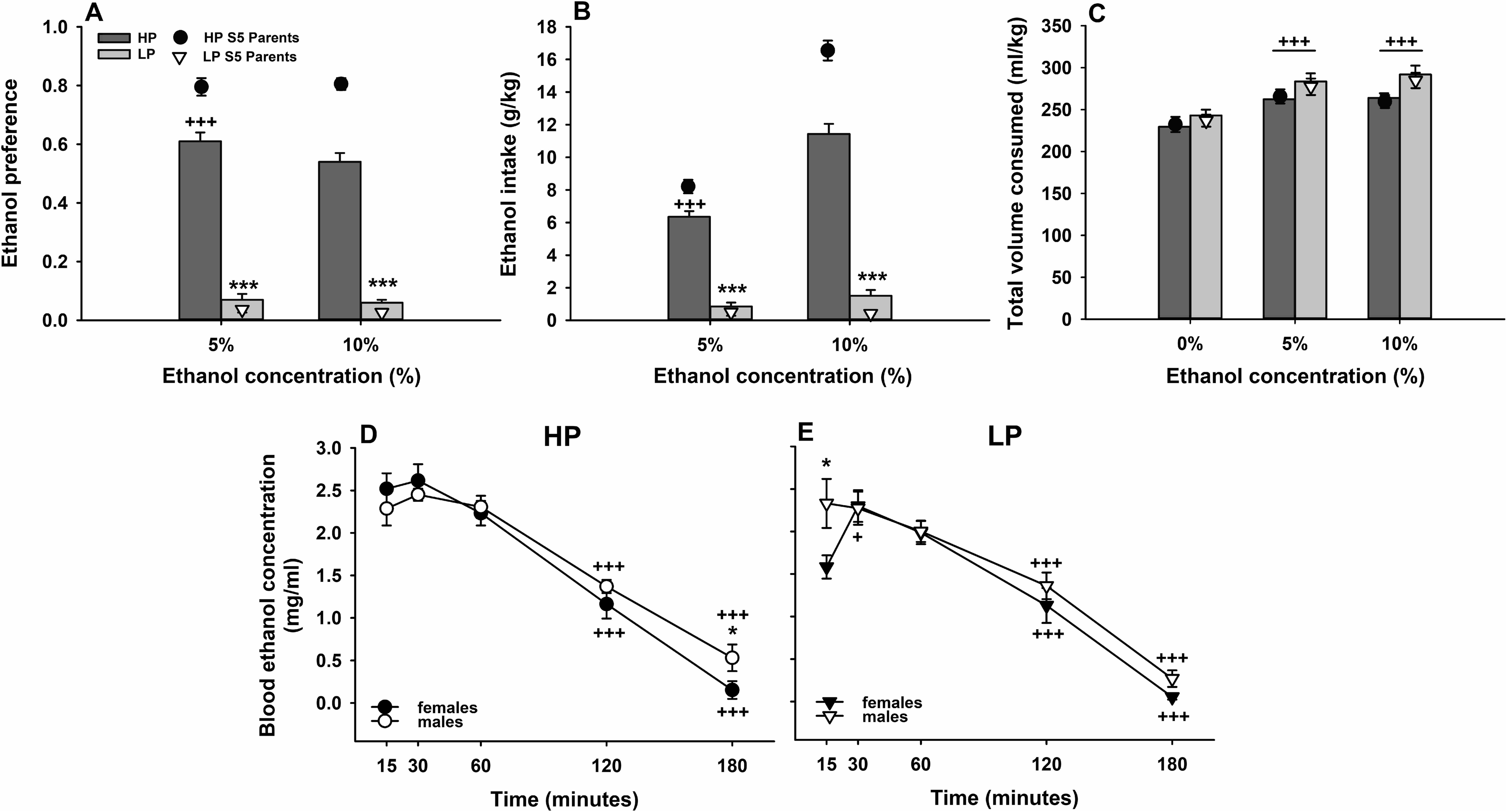
Ethanol Intake, Preference and Clearance: ***(A-C)*** Ethanol preference, ethanol intake and total volume consumed for S4 generation high preference (HP) and low preference (LP) mice. Shown are means ± SEM for (A) ethanol preference (ml from the ethanol tube / total ml); (B) ethanol intake (g/kg); and (C) total volume consumed (ml/kg) from the water and ethanol tubes. Also shown in each panel are means ± SEM for the S4 offspring chosen based on ethanol preference to serve as the S5 breeders to produce mice for additional behaviors, and for metabolism and transcriptome studies. N=116 HP (62 female and 54 male) and 67 LP (37 female and 30 male) mice; mean age (± SEM) when ethanol was first offered 67 ± 0.2 days (range = 56-70 days). HP S5 Parents, n=24 breeders per sex; LP S5 Parents, n=24 breeders per sex. ***p<0.001 for the effect of line at the indicated concentration; +++p<0.001 for the difference from 10% within HP or LP line (A and B) or from 0% (C). (D-E) Shown are means ± SEM for blood ethanol concentrations in blood samples obtained at 15-180 min after 2 g/kg IP ethanol injection in (D) HP and (E) LP female and male mice from the S4 generation. Each mouse contributed a 20 µl lateral tail vein sample at each time point. N=10-11/line/sex; mean age (± SEM) was 97 ± 0.4 days (range = 90-101 days). *p<0.05 for the effect of sex; +p<0.05, +++p<0.001 compared to T15.

#### 2.1.2 Ethanol intake

Data are shown in Figure 1B for ethanol intake (g/kg). There was a significant main effect of sex (F[1,167]=7.1, p<0.01), but no interaction of sex with line or concentration. Females consumed an average of 5.8 ± 0.4 g/kg, whereas males consumed an average of 4.2 ± 0.4 g/kg ethanol (collapsed on concentration). There was a significant line by ethanol concentration interaction (F[1,167]=40.5, p<0.0001). Simple main effects analysis identified a significant line difference for both ethanol concentrations, with the HP mice consuming more ethanol than the LP mice (ps<0.001). There was no impact of ethanol concentration on intake within the LP mice; however, the HP mice consumed more ethanol when it was offered as a 10% concentration (p<0.001). Mean ± SEM ethanol intake (g/kg) values for the 5 and 10% ethanol concentrations are also shown for the subset of S4 mice chosen to serve as S5 breeders (based on ethanol preference).

#### 2.1.3 Total volume consumed

Data for total volume consumed (ml/kg) are shown for the water only period (0%) and during the time when 5 and 10% ethanol were offered (see Figure 1C). Repeated measures ANOVA, with concentration (0, 5 and 10%) as the repeated measure, and selected line and sex as independent factors, identified significant main effects of sex (F[1,165]=15.2, p<0.001) and concentration (F[2,330]=63.0, p<0.001). Sex interacted with concentration (F(2,330)=4.2, p<0.05) but not line, so data in Figure 1C are shown for the lines collapsed on sex. Females consumed more total fluid in ml/kg during all periods (ps<0.05-0.001) and both lines consumed more total fluid during the times when one bottle contained ethanol than when both contained water (ps<0.001). The means ± SEM were 246.3 ± 6.7, 291.7 ± 6.9, and 289.2 ± 6.5 ml/kg for females for 0, 5, and 10% ethanol, respectively. For males the means ± SEM were 220.9 ± 6.8, 245.0 ± 7.4, and 256.5 ± 8.3 ml/kg for 0, 5 and 10% ethanol, respectively. Mean ± SEM total volume consumed for the S5 breeders is also shown in Figure 1C.

#### 2.1.4 Correlations

Ethanol preference and g/kg ethanol intake were highly correlated. For 10% ethanol, the correlation in the S4 offspring was r=0.94 (p<0.001; N=174). In comparison, the correlation between ethanol preference and total volume (ml/kg) during the time that 10% ethanol was offered was r=-0.17 (p<0.05), and between ethanol intake and total volume, it was r=0.04 (NS), indicating little relationship. Correlational data are graphed in the Supplemental Information (Figure S1).

#### 2.1.5 Blood Ethanol Elimination Rate

Blood ethanol concentration (mg/ml) data across time were examined by ANOVA, with time as a repeated measure, and selected line and sex as independent factors. There was a significant line by sex by time interaction (F[4,128]=4.1, p<0.01), therefore, data were examined for sex differences within each line. For both HP and LP mice, there was a significant sex x time interaction (F[4,64]=2.7, p<0.05 and [F4,64]=2.9, p<0.05, respectively). Follow-up simple main effect analyses identified a sex difference only at T180 (p<0.05) in HP mice and T15 (p<0.05) in LP mice, with lower BEC in females than males in both cases. As can be observed in Figure 1D and 1E, peak BECs occurred at 15-30 minutes post treatment and reached near 0 values at 180 minutes. Because sex differences occurred at isolated times, we considered whether the lines exhibited different patterns of ethanol elimination when data for the sexes were combined and found no evidence for a line x time interaction (p=0.67). Rate of elimination was 0.96 and 0.92 mg/ml/h for HP and LP mice, respectively.

#### 2.1.6 Other Behavioral Traits

Detailed results for additional phenotypes measured in the HP and LP mice are presented in the Supplemental information. Preference values for saccharin and sucrose were high in both selected lines (Figures S2A and S3A), whereas quinine was avoided, particularly at the higher concentration (Figure S4A). The only significant line difference for preference was when the lower concentration of saccharin was offered (HP > LP). For consumption (mg/kg or g/kg), compared to LP mice, HP mice consumed more of both sweet solutions (Figures S2B and S3B), but the selected lines did not differ in consumption of the bitter tastant (Figure S4B). Ethanol induced locomotor stimulation in both lines, but to a greater extent in LP than HP mice (Figure S5). BEC at the end of the locomotor test was dose-dependent but did not differ between the lines (Figure S5A inset). Finally, ethanol induced time- and dose-dependent hypothermia, but there was no difference in hypothermic response between the HP and LP lines (Figure S6).

### 2.2 Single-Nuclei RNA Sequencing

We collected and analyzed snRNA-Seq data to identify and characterize the effects of bidirectional selection for ethanol preference on individual cell-type populations in the CeA using tissue from 14 adult mice (7 HP, 7 LP; n = 3-4/sex) which passed filtering during coarse clustering (see Section 4.4.2). We note no significant difference (false discovery rate (FDR) < 0.05) was detected between the HP and LP animals in inhibitory neurons or non-neuronal cells regardless of the analytic strategy (either differences in cell-type composition or expression). The only exception is 6 genes that were significantly differentially expressed (DE) across excitatory neuronal clusters. Given the variance across individual samples, we assume the power to detect a significant selection difference was not sufficient. Cell-type populations identified here were used to extend the analysis of higher-powered bulk RNA-Seq data in Sections 2.3 and 2.4. Markers for all identified cell-types discussed below are available in the Supplemental information and on GitHub. Compositional results are given in Supplemental Table S1, while summarized DE results for the excitatory cells are given in Supplemental Table S2.

#### 2.2.1 Overall Clustering

An initial clustering of all 46,219 cells that passed filtering found 39 clusters, which were robustly identified as either neuronal or non-neuronal based on their expression of previously identified markers (see Section 4.4.3 for details, and Supplemental Table S3). The neuronal cells were separated into discrete populations as either inhibitory (17 clusters with 22,221 cells) or excitatory (11 clusters with 10,287 cells), while the non-neuronal clusters (11 clusters with 13,731 cells) corresponded to the expected populations of microglia, astrocytes, oligodendrocytes and their precursors, endothelial cells and a small population of immature neurons. Each subpopulation of cells (inhibitory, excitatory and non-neuronal) was then re-analyzed (normalized, clustered, etc.) separately to maximize discriminatory power of cell-type markers within similar cell types.

No significant differences in expression nor composition were found between the HP and LP mice in the broad categories (inhibitory, excitatory or non-neuronal). Coarse clustering results are shown in Figure 2, where 2A is a UMAP of all cells that passed filtering, identified as shades of blue, red and green for inhibitory, excitatory and non-neuronal cells, respectively. Figure 2B shows the relative expression of marker genes across the three major cell types. Figure 2C shows the percentage of cells for each sample that were identified as inhibitory (∼48%), excitatory (∼23%) or non-neuronal (∼29%), showing no difference between HP and LP mice. We note that the CeA shows a roughly 2:1 ratio of inhibitory to excitatory neurons. Additionally, non-neuronal cell populations are generally discarded during analysis of snRNA-Seq data in the CeA either explicitly or by filtering “low” UMI/count cells, but we included this significant population (making up ∼29% of all cells) in our analysis as these cells may explain phenotypic differences between the HP and LP mice [22–25].

**Figure 2.**
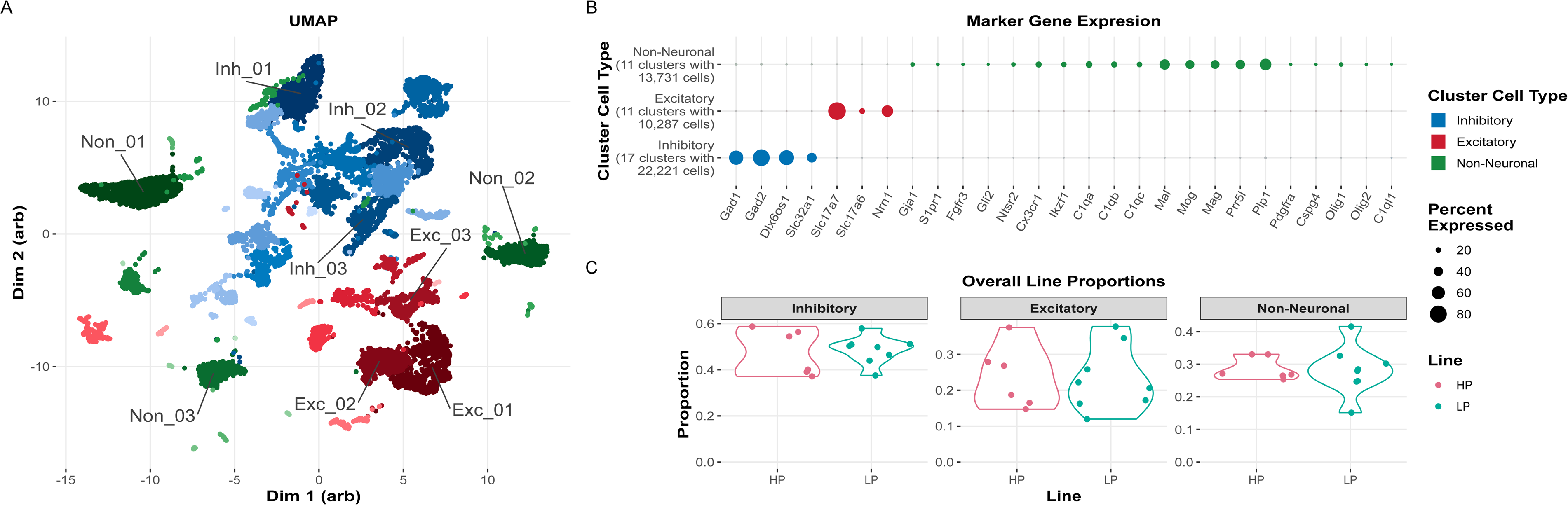
Overall Clustering: Clustering of all 46,219 cells that passed filtering, showing a robust separation of inhibitory neurons (shades of blue), excitatory neurons (shades of red), and non-neuronal cells (shades of green) in clusters (A) and by marker gene expression (B). No differences in cell-type compositions were observed between the HP and LP lines (C), but inhibitory neurons (48%) were twice as common as both excitatory neurons (23%) and non-neuronal (29%) cells.

#### 2.2.2 Inhibitory Neurons

Overall clustering and filtering identified 22,221 inhibitory neuronal cells based on their expression of *Gad1, Gad2, Slc32a1* and *Dlx6os1* and the lack of expression of other cell-type specific markers (see Section 4.4.3). These inhibitory neurons represented ∼49% of all cells and demonstrated great diversity by clustering into 18 final, distinct clusters after the processing described in 4.4.3. One cluster with fewer than 100 cells was discarded, leaving 22,123 cells. Inhibitory neurons were divided into two main branches based on their DE of *Wfs1*, *Meis2, Zfhx3*, and *Zfhx4* (Inh 1-6, 8, 10-11, 13-14, 18) versus *Maf* and *Zeb2* (Inh 7, 9, 12, 15-17). This organization indicated an expected split between projecting neurons versus local interneuron populations.

These clusters, as well as their markers, are visualized in Figure 3. Figure 3B is a UMAP showing the isolation of the 18 clusters. Figure 3A is a dotplot showing the relative expression of marker genes that differentiate an inhibitory neuronal subtype from the remaining inhibitory neurons, and here the subtypes have been ordered according to their similarity based on an unsupervised hierarchical clustering on the Euclidean distance of their highly variant genes. A dendrogram of this clustering is provided for visual clarity in Figure 3A, and more fully in Figure 3C where genes that separate onto different sides of each split (left: green, right: blue) are also included. We remind the reader that in general, a positive marker on one side is a negative marker on the other, but the opposite is not always true. Additional details on the identification of dendrogram markers is available in the Methods (Section 4.4.3) within the discussion on clustering. For example, as one moves from left to right (from Inh_18 towards Inh_8) on the left major branch (*Meis2, Wfs1, Foxp2* expressing) one observes a diversification of receptor types away from dopamine (*Drd1* and *Drd2*) towards others (Glycine [*Glra3*], Glutamate [*Grik4, Grin3a]*). One may also examine what differentiated similar inhibitory neuron subtypes, such as Inh_3 and Inh_4, which expressed different dopamine receptors (*Drd1* vs *Drd2*). As expected, cell types are finely differentiated by their expression of neurotransmitters and receptors and coarsely differentiated by translation factors.

**Figure 3.**
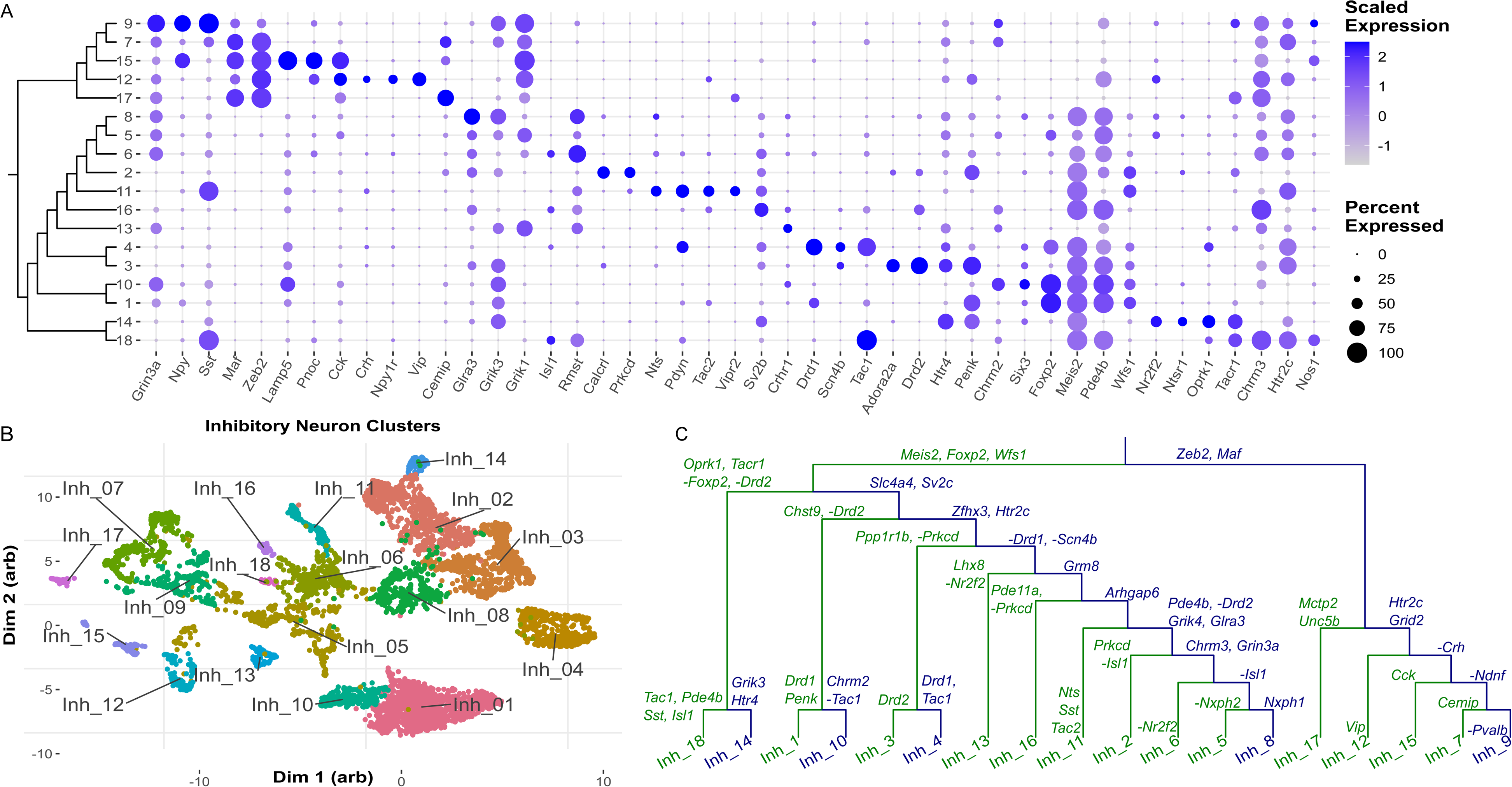
Inhibitory neuron clustering: Analysis of 22,221 inhibitory neurons identified 18 clusters corresponding to cell populations with distinct expression profiles. (A) Marker gene expression is shown for each of the 18 clusters, and clusters are ordered according to their similarity as determined by hierarchical clustering on their gene expression. (B) Clusters are shown visually in a UMAP. (C) A dendrogram of cluster similarity, where at every split of the dendrogram unique markers are identified for the left (green) and right (blue) branches. The major split isolates *Meis2*, *Foxp2* and *Wfs1* expressing inhibitory neurons from those expressing *Zeb2* and *Maf*.

As with the overall clustering, no differences in expression or composition were identified between the HP and LP lines across these 18 inhibitory clusters. Composition is visualized in Supplemental Figure S7.

#### 2.2.3 Excitatory Neurons

Roughly half as many excitatory neurons were identified from the overall clustering, with 10,287 cells used for finer clustering, representing ∼23% of all cells that passed initial filtering. Excitatory neurons were identified by their expression of *Vglut1 (Slc17a7), Vglut2 (Slc17a6)* or *Nrn1* and the lack of expression of other cell-type specific markers (see Section 4.4.3). Fine clustering identified two clusters dominated (>80%) by a single sample, which were then excluded, leaving 9740 cells. Ten excitatory neuronal subtypes were identified and may be broadly classed by their expression or *Vglut1* and *Vglut2*. All but one cluster expressed *Vglut1*, with cluster Exc_10 being the lone cluster that expressed only *Vglut2*. Three other clusters, Exc_6, Exc_9 and Exc_5 were found to express both *Vglut1* and *Vglut2*.

These clusters and their markers are visualized in Figure 4 in the same way as the inhibitory clusters were shown in Figure 3. Whether a cluster expressed *Vglut1* or *Vglut2* is additionally indicated at the bottom of the dendrogram in Figure 4C. We note that Exc_10 is the only cluster that did not express *Gabra1* and *Gabrb2*, and the only cluster to express *Nts* and *Tacr1.* Exc_10 is also the smallest identified cluster, representing roughly 2% of cells (see Supplemental Figure S7).

**Figure 4.**
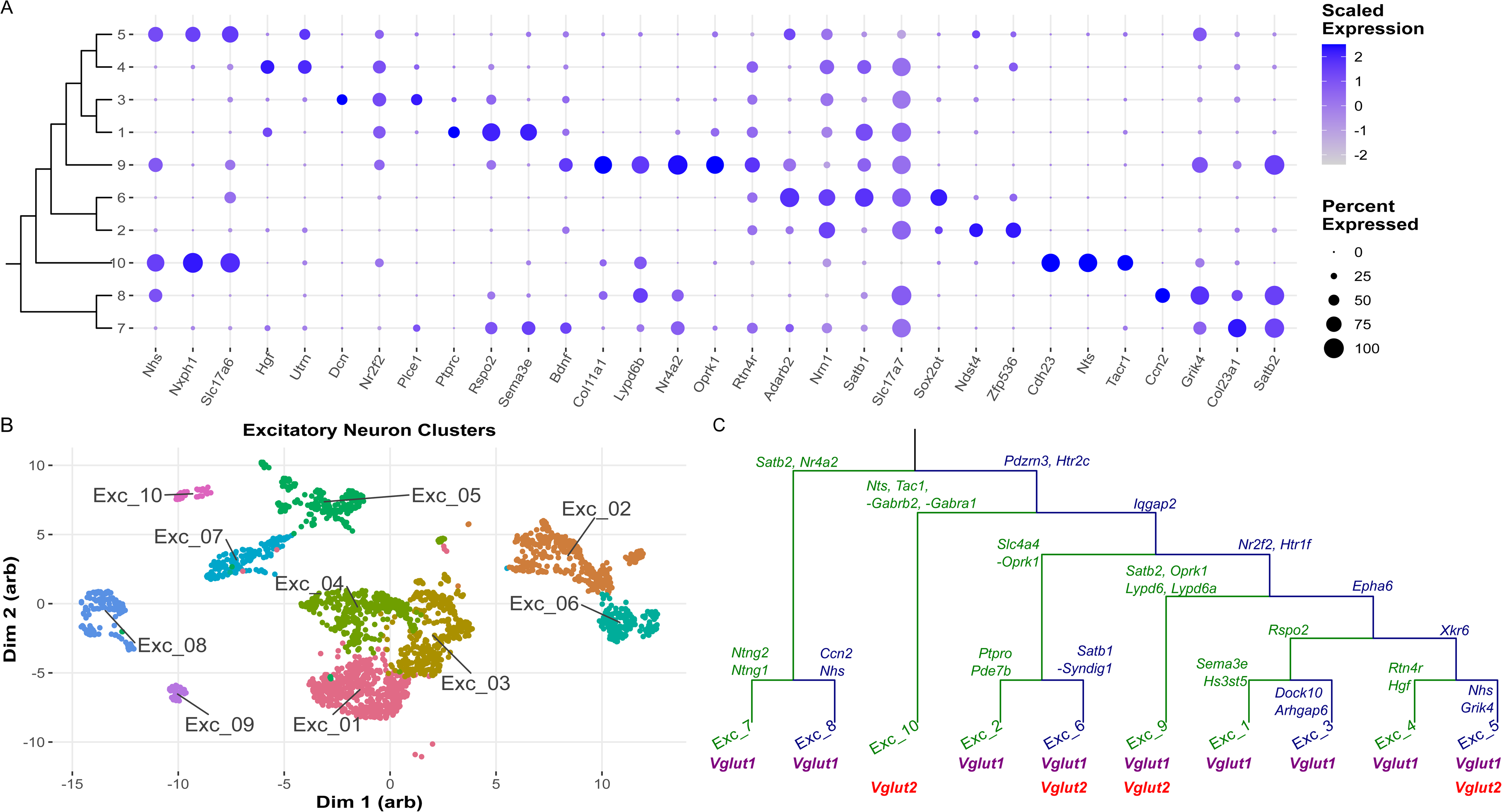
Excitatory neuron clustering: Analysis of 10,287 excitatory neurons identified 10 clusters corresponding to cell populations with distinct expression profiles. (A) Marker gene expression is shown for each of the 10 clusters, and clusters are ordered according to their similarity as determined by hierarchical clustering on their gene expression. (B) Clusters are shown visually in a UMAP. (C) A dendrogram of cluster similarity, where at every split of the dendrogram unique markers are identified for the left (green) and right (blue) branches. Expression of *Vglut1* (*Slc17a7*) and *Vglut2* (*Slc17a6*) is indicated below each cluster.

Many excitatory clusters express *Rspo2* or *Satb1,* identified as markers associated with the basolateral amygdala (BLA) by Hochgerner et al. [22]. The authors find it unlikely that the same amount of BLA would be incidentally dissected across 14 samples, However, we acknowledge that, given the small number of samples and the known individual variation of the HP and LP lines, it is not possible to determine whether these excitatory neurons are associated with the BLA or found within the CeA of the HS-CC mice, as comparisons can only be made to studies on B6 backgrounds, and full cell proportions of non-inhibitory neurons are not typically reported. Additionally, the BLA has projections to the CeA [26], so it is also possible that these genes were expressed in projection neurons terminating in the CeA.

No differences in composition (visualized in Supplemental Figure S7) and few differences in expression were identified between the HP and LP lines across these 10 inhibitory clusters. A total of 6 genes were found to be DE in the excitatory clusters: Exc_9 (*Cck, Cdh8, Nrxn3, Pex5l*), Exc_8 (*Fign*), and Exc_6 (*ENSMUSG00000095041*). The pseudo-expression of these genes is shown in Supplemental Figure S8, where it is apparent that all differentially expressed genes (DEGs) in Exc_9 were driven by outliers. *Fign* was down expressed in the HP mice and was also found to be differentially wired (DW, Section 4.4.5) and differentially variable (DV, Section 4.4.5) in the bulk RNA-Seq results (Section 2.3). *ENSMUSG00000095041* was up expressed. Summarized DE results are available in Supplemental Table S2, whereas full results are available at GitHub.

#### 2.2.4 Non-Neuronal Cells

Fine clustering of the 13,731 non-neuronal cells identified 3 clusters with membership below 10 cells average per sample, and two doublet clusters. Removing these clusters resulted in 12,454 cells remaining that robustly clustered into four clusters corresponding to the expected populations of oligodendrocytes and their precursors, microglia, and astrocytes. No differences in composition or expression were identified between the HP and LP lines, visualized in Supplemental Figure S7. These clusters, as well as their markers, are visualized in Figure 5 in the same way as the inhibitory clusters were shown in Figures 3 and 4; however, no dendrogram is presented.

**Figure 5.**
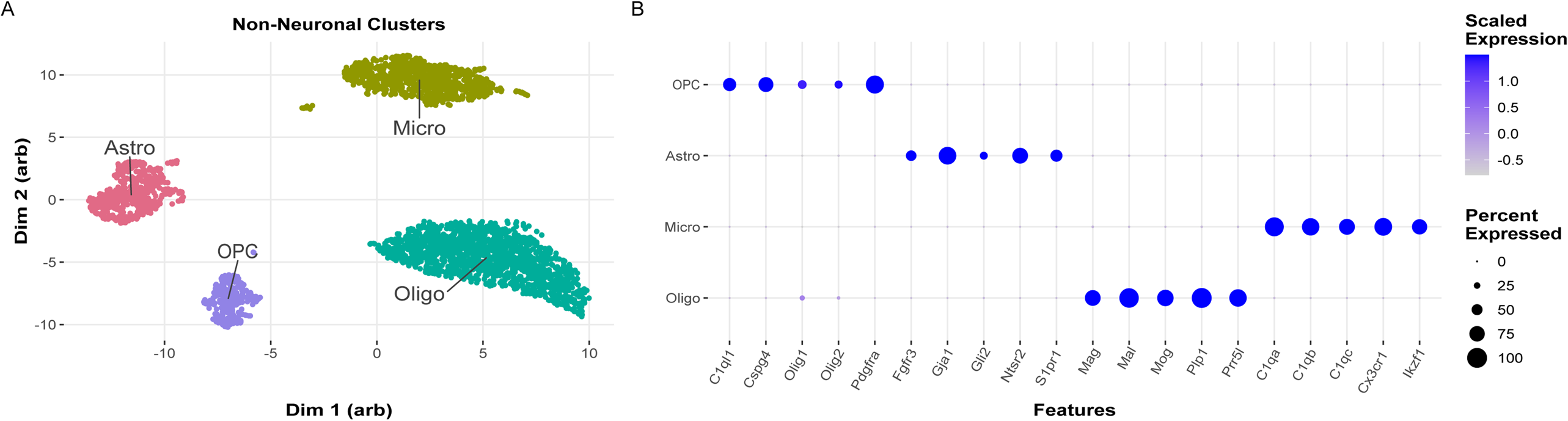
Non-neuronal clustering: Analysis of 13,731 non-neuronal cells identified 4 clusters corresponding to unique populations of oligodendrocytes and their precursors, microglia and astrocytes. (A) Clusters are shown visually in a UMAP. (B) Marker gene expression is shown for each of the clusters, which are labeled according to their expression of canonical marker genes.

### 2.3 Bulk RNA Sequencing

HP and LP CeA samples, n = 100 per line balanced for sex, were analyzed; 20 million paired-end reads were obtained per sample. RNA-Seq data for seven samples did not pass our Q/C metrics (see Methods and Supplemental Table S4); the deleted data were balanced across line and sex. 16,198 genes and gene-like features with ≥ 0.5 counts per million reads met the threshold for further analysis; 14,287 were protein coding (Supplemental Table S5).

#### 2.3.1 Principal Component Analysis

To assess data quality and signal-to-noise ratio for categorical variables of biological interest (sex and line), compared to technical effects (e.g. sequencing batch), dimensionality reduction was carried out using principal component analysis (PCA). PCA was performed via the covariance matrix of all 193 samples and the 16,198 genes that passed filtering (additional details are in Supplemental Table S6). In total, sex and line were significantly associated with 21.2% of the explained variance (PCs 2-5 and 8) and batch was significantly associated with 8.8% of the explained variance (PCs 6, 7 and 12). This is a signal-to-technical noise ratio of 2.4x, which is in line with expectation given a short-term selection from a heterogeneous stock and indicated the expression data were of sufficiently high quality for downstream analysis.

The PCA data are illustrated in Supplemental Figures S9A and S9B where PC scores of individual mice are plotted in green (LP), brown (HP) and as solid circles (male) and opaque triangles (female). Supplemental Figure S9A shows PC3 [significantly associated with line, FDR = 4.4e-6] vs PC2 [significantly associated with sex, FDR = 4.6e-3]. Margin plots show the data grouped by line (PC3) and sex (PC4) in box plots, for which median separation is more apparent. Figure S9B shows PC5 vs PC4, which are both significantly associated with sex and line [FDR < 3e-40 for a combined Sex and Line factor]. Margin box plots show data grouped by sex and line. It is apparent that PCs 4 and 5 separate samples across line and sex, with female LP mice clustering away from male HP mice along PC5 and male LP mice clustering apart from female HP mice along PC4. Given that PCs 4, 5, 6 and 8 (9.8% of explained variance) were associated with multiple factors, all PC scores were tested using a 3-way ANOVA to determine if any significant interactions were present. No significant interactions were identified.

#### 2.3.2 Bulk Differential Gene Sets

Data were extracted for DE, DV and DW. Additional details on DV and DW are in Colville et al. [7, 8] and Section 4.4.5. Because sex and line each explained significant amounts of variance in the expression data, differential analyses were carried out using sex and line explicitly in the model. No sex by line interactions for differential genes were identified, thus, contrasts were constructed collapsed on sex to focus results on selected line differences; main effects of sex will be considered in a future work. (Note that an FDR < 0.05 was the threshold for significance for all statistical tests). We began by investigation of the gene ontological (GO) enrichment of these gene sets. This provides a broad and relatively low-level understanding of the transcriptional changes driven by preference selection.

The DE, DV and DW differences between the HP and LP lines are summarized in Figure 6A; the full results are found in Supplemental Tables S7, S8 and S9. 2996 genes were DE, roughly equally divided between those genes that had higher and lower expression in the HP line. Some of these genes include *Gabra2*, ***Gabrb1,*** *Gabrd, **Gabrg1**, **Gabrq**, **Glra1, Glra3**, Grik3, Grin2a, Grin2b, Htra1, Htra2, Htr2a, **Htr7**, Npy, **Npy1r, Npy5r*** and *14 protocadherin genes.* (Genes in bold had lower expression in the HP line.) The focus on this subset of DEGs, especially the GABAergic and glutamatergic genes, aligns with the analysis below on co-expression networks and the single nuclei data. 426 genes were significantly DV; here the data were 10:1 in favor of lower variability in the HP line (Figure 6A). Some of the DV genes include ***Adora1, Atf6, Chrm4, Chrnb2, Gabra2, Glra2, Glra3, Htr1d, Htr2c, Nts, Ntsr1*** and ***Ntr2k.*** The difference in variance for ***Gabra2*** between the lines is illustrated in Figure 6B. 407 genes were significantly DW; 70% had a decrease in correlation. The median number of edges affected was relatively small (3/gene) although for some genes it was substantially higher; e.g. 70 for *Cdc7*. Some of the DW genes include ***Arc, Atf6, Cck, Ephb6, Gabre, Gabrq,*** *Htr1d, **Htr2c,** Lmo3, **Oprm1, Oxtr,*** and ***5 collagens.*** *Oxtr* (oxytocin receptor) had 23 edges that included *Alk* [27]. A heat map illustrating the DW for *Ephb6* (ephrin receptor type b6) is illustrated in Figure 6C. Of interest, significant edge changes were observed for *Nts* and *Oprm1*. Figure 6D illustrates the enriched ontologies for the DE, DV and DW datasets; for each group, a significant enrichment in receptor activity or synaptic transmission was detected (additional ontology details are found in Supplemental Table S10).

**Figure 6.**
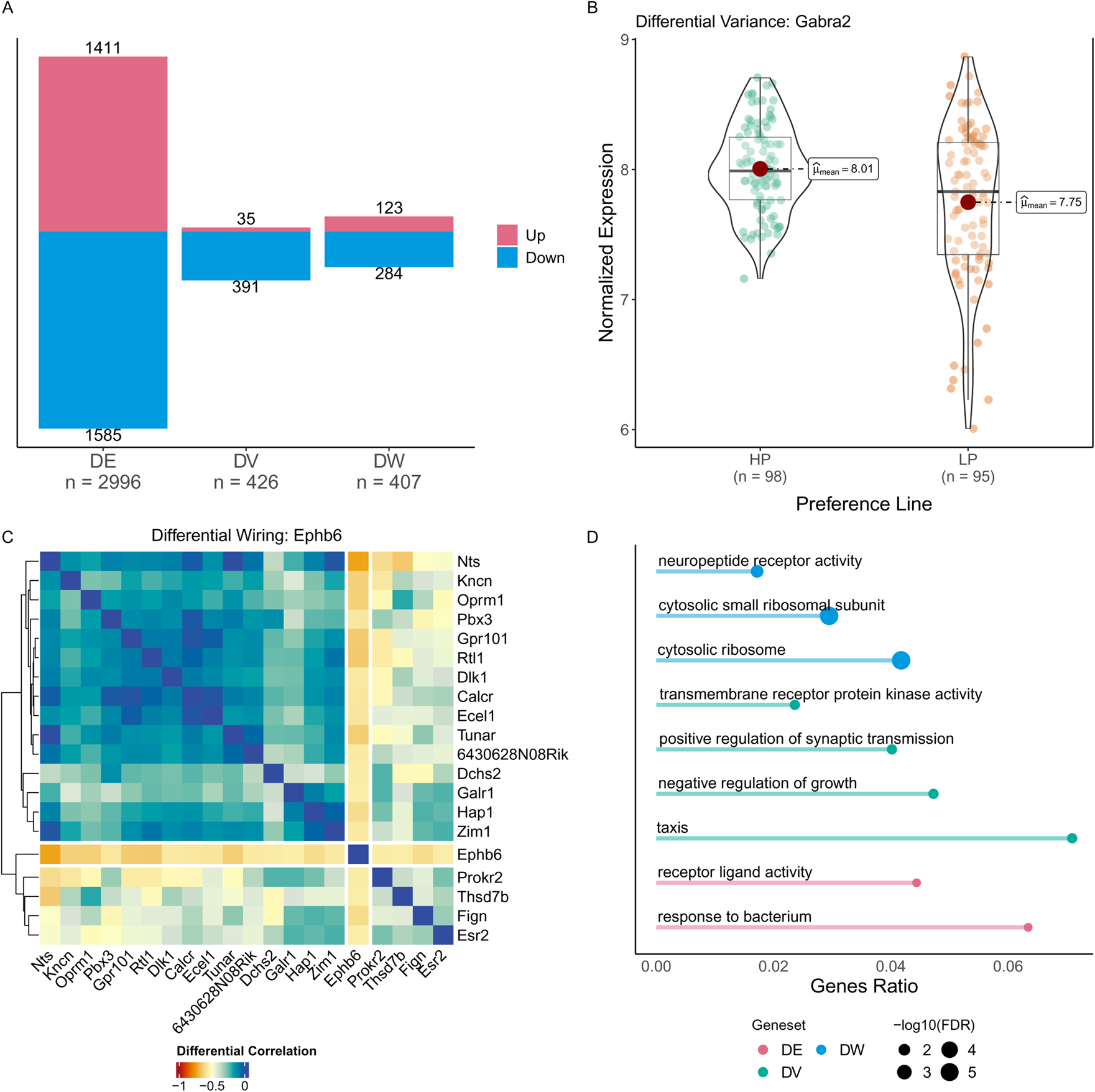
Differential Genes Overview: (A) The number of Differentially Expressed (DE), Differentially Variable (DV) and Differentially Wired (DW) genes detected in the CeA for HP vs LP mice. Up genes are shown in red, where ‘up” means higher median expression in the HP mice for DE genes, higher variance in the HP mice for DV genes, and a higher total correlation across differential edges in the HP mice compared to the LP mice for DW genes. Examples of a (B): differentially variable and (C): differentially wired gene. (B) *Gabra2* is a differentially variable gene, where variance is significantly lower in the HP mice as compared to the LP mice using an F-test. *Gabra2* is also differentially expressed. (C) *Ephb6* is a down differentially wired gene that has 35 differential edges, defined as an absolute change in gene-by-gene correlation of greater than 0.5 between the HP and LP lines. Shown here as a heatmap of the change in absolute gene-by-gene correlation from the HP to LP lines for *Ephb6* and its top 20 differential edges. Differential edges are shown in yellow to orange, grey indicates an absolute change in gene-gene correlation slightly below the threshold of 0.5, while green to blue indicates lower absolute changes. We note that *Prokr2*, *Thsd7b*, *Esr2*, and *Fign* show a similar change pattern in their gene-by-gene correlation to *Ephb6*, while the remaining genes do not. This gene is shown as it is differentially edged to many hub genes in the GABAergic weighted co-expression network module to be discussed later. (D): Ontological enrichment of DE (red), DV (green) and DW (blue) genes (top 750 by abs(logFC)). The length of the line indicates the ratio of DE/DV/DW genes within the GO gene-set, and the size of the circle indicates significance (-log10). All significantly enriched GO terms are shown to DE and DW genes, but only 4 of 13 significantly enriched GO terms are shown for DV. The four shown are the most different from one another in terms of their semantic similarity (gene overlap), to maximize the information shown.

### 2.4 Weighted Gene Co-Expression Network Analysis

As in Colville et al. [7, 8], we used WGCNA to investigate and annotate the bulk gene expression data. However, rather than use a consensus module approach to analyze the LP and HP populations, the HP and LP expression networks were formed independently. Note the modules in the HP network are labelled numerically and those in the LP network alphabetically. How the modules are labelled has no biological significance. These data are summarized in Supplemental Table S11. Supplemental Table S11 also denotes the hub genes in each module and the unique hub genes found only in the HP or LP networks. There were ∼500 unique hubs in both the HP and LP networks or approximately 25% of the total hub genes per network. A similar number of modules was identified in each network with 20 found in the HP network, and 21 in the LP network.

Supplemental Table S12 summarizes the strategy used to characterize the ontology of network modules, where module membership was tested for enrichment against gene-sets of interest, including differential genes, snRNA-Seq cell-type markers, and risk signatures identified in the HS-CC founders. Of the 8 inbred strains used to form the HS-CC, two of the strains (B6 and PWK) have high ethanol preference [16]. Anderson et al. [17] compared gene expression in the B6 and PWK strains to gene expression in the 6 non-preferring HS-CC strains. Genes showing a difference in expression between the preferring and non-preferring strains in the same direction were noted as ‘signature genes’. The list of the B6 and PWK signature genes, likely to be ones impacted by selection, and the modules in which they appear are found in Supplemental Tables S13 and S14. The number of DE, DV and DW genes within each module is given along with total genes per module. As noted above, DE, DV and DW up and down refers to the changes in expression when comparing the HP to the LP genes. Supplemental Tables S7, S8 and S9 list the differential genes and the modules in which they appear. Note that not all differential genes and signature genes are found in network modules.

Gene ontologies for each network module are found in the Supplemental data HP and LP module ontology tables, and on the GitHub. This analysis revealed that 11 of the HP modules (3, 6, 7, 8, 9, 10, 11, 14, 15, 16, and 17) and 9 of the LP modules (B, F, I, J, K, M, O, P and Q) were enriched in genes associated with synaptic function. As noted above, the single nuclei data were used to create a CeA ontology that focused on subtypes of inhibitory and excitatory neurons. Application of these ontologies to the network modules is found in Supplemental Table S6. A schematic of Supplemental Table S6 is provided in Figure 7 where the enrichment of modules with gene sets of interest is visualized. Only modules that were enriched in gene-sets associated with selection (differential genes, unique hubs) and whose hub genes showed ontological enrichment are shown. We were most interested in modules that: (1) were associated with selection, (2) were preserved between both networks and (3) had enrichment in cell-type specific markers.

**Figure 7.**
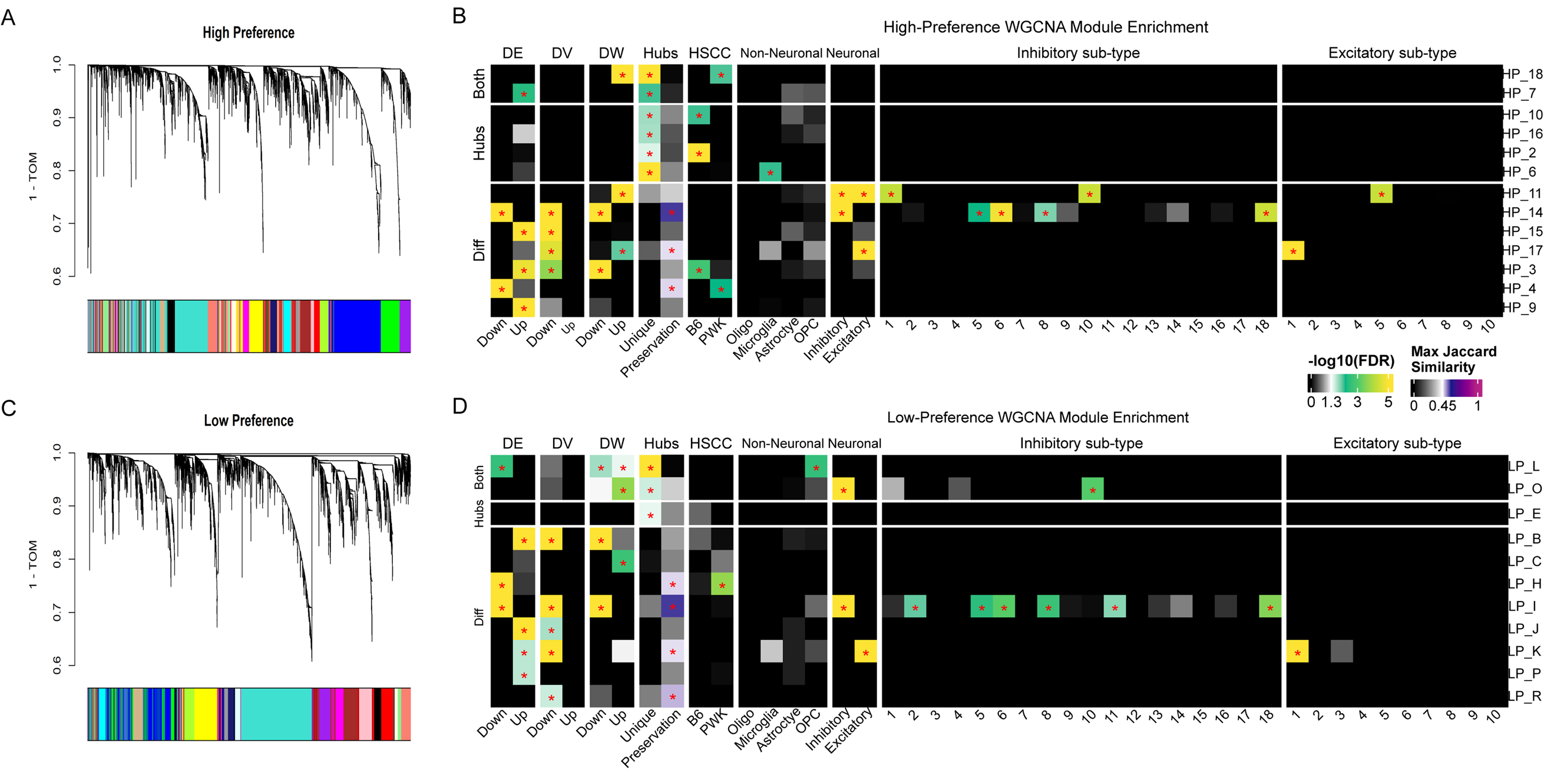
WGCNA Overview: An overview of weighted gene co-expression network analysis (WGCNA) results for the high preference (HP) and low preference (LP) lines. The top row (A and B) shows results for the HP network, whereas the bottom row (C and D) show results for the LP network. The left columns (A and C) show dendrograms resulting from hierarchical clustering of the HP (left) and LP (right) genes into modules based on a dissimilarity measure of 1 – TOM (topological overlap network). The right columns (B and D) are heatmaps showing the enrichment of the HP (top) and LP (bottom) modules by various gene sets of interest. Any shading from white to green to yellow (also marked with a red asterisk) indicates significant enrichment at FDR < 0.5. Neuronal sub-types are only considered if the corresponding overall neuron (e.g. “Inhibitory”) is enriched. If a module is preserved (has a Jaccard similarity above 0.47), then the preservation column is shaded from white to purple. In all cases, grey to white indicates insignificant enrichment, or a lack of preservation. Column groups are, from left to right: (1-3) Differential genes (up or down in HP, for expression, variability and wiring), (4) Hubs unique to that network, (5) HS-CC founder risk signatures for B6 and PWK, (6) Non-neuronal cell markers, (7) Neuronal cell markers, (8) Inhibitory sub-type markers, and (9) Excitatory sub-type markers. Rows are organized by whether a given module is enriched in Differential genes, Hub genes, or both.

The following summarizes key points extracted from the network analyses.

1. ***The HP and LP modules were more distinct than similar.*** There are seven HP/LP pairs where the modules appear significantly similar [Jaccard Similarity (JS) ≥ 0.45; see Section 4.4.5], or roughly 35% of all modules. By evaluating their ontological enrichment, we found that three of these pairs (HP_8/LP_F, HP_14/LP_I and HP_17/LP_K) involve modules with synaptic annotations. The HP_4/LP_H and HP_13/LP_R pairs are involved in RNA processing and the HP_5/LP_D pair is involved in ciliary function, here assumed to be primary cilia function. The final pair, HP_1/LP_A, is involved in myelin. *Note, Only the HP_14/LP_I and HP_17/LP_K pairs satisfy all three of the criteria above*.
2. ***The LP network was consistently more connected than the HP network.*** kWithin and kOut, the intra- and inter-module connectivity, are (in sum) for all modules larger in the LP line as compared to the HP line. These data are consistent with the decrease in variability in the HP line (see Figure 4). Focusing on the three synaptic pairings noted above in “1”, the HP/LP ratios for the mean kWithin are 0.82, 0.28 and 0.88, respectively. For just the hub nodes (see bottom panels of Supplemental Table S6), the ratios are 0.91, 0.32 and 1.00, respectively. The difference between the HP_14 and LP_I modules can also be seen in measures of density, centralization and heterogeneity [21]. The HP module has ∼2x lower density (lower connectivity among members of the module), has ∼2x lower centralization (hub nodes are less important) and is 20% more heterogeneous.
3. ***The HP_14/LP_I* modules were enriched in genes down expressed in the HP mice, and inhibitory neuronal subtypes.** For the HP_14/LP_I comparison, the kWithin differences for the top 25 hub nodes in each module are illustrated in Figure 8 and show a marked loss of connectivity in the HP_14 module compared to the LP_1 module. Nineteen of the top 25 HP hub nodes are found in the top LP hub nodes. Both the HP_14 and LP_I modules were significantly enriched in DE, DV and DW genes; note that in both modules, the enrichments were down (HP<LP). Neither module was enriched in B6 or PWK signature genes although overall the HP modules were more enriched in these signature genes than the LP modules (8 vs 4). Both the HP_14 and LP_I modules were enriched in genes associated with two inhibitory neuronal sub-types characterized by their *Isl1* and *Tac1* expression: Inh_6 and Inh_18.Cluster Inh_6 was additionally enriched in the following genes: *Tshz2, Tacr3, Chrm3, Gaba5, Gabrg1* and *Nts,* identified as being key to CeA function and/or associated with one or more ethanol phenotypes (e.g. see [23, 28]). Cluster Inh_18 was enriched in the following genes: *Dgkk. Sst. Nos1, Chrm3. Tacr1,* and *Grik4* but not *Nts.* HP_14 and LP_I were also enriched in markers from Inh_5 and Inh_8, two subtypes that were closely related to Inh_6 in terms of their overall expression patterns (See Figure 3C), but are distinguished by their expression of *Grik3, Nphx2* or *Nphx1,* and lack of *Isl1*. All 5 of these inhibitory neuronal clusters express the hub genes *Baiap3* and *Gpr101*. The LP_I module is additionally enriched in Inh_2 and Inh_11 markers, which cluster along with Inh_5, 6 and 8, but are differentiated by expression of *Prkcd* (Inh_2) or *Nts, Sst* and *Tac2* (Inh_11).
4. **The HP_17/LP_K pairing was enriched in genes up expressed in the HP mice, and excitatory neuronal subtypes.** The within module connectivity schematics for the top 25 hub nodes in each module are illustrated in Figure 9; in contrast to the HP_14/LP_I differences, the kWithin differences here are small. Thirteen of the top 25 HP nodes are found in the top 25 LP nodes; however, the overall ontologies for the 2 modules, determined by all the hub nodes (top 20% of all nodes) are very similar (see Supplemental Ontology Tables). A notable exception was the LP enrichment in hormone receptor binding; genes in this category include *Cck, Vip* and *Jak1.* Related to this ontology, both modules HP_17 and LP_K were enriched in genes from the most common excitatory subtype Exc_1 (see Figure 4), which expressed only *Vglut1* and is differentiated from other excitatory neurons by its expression of *Sema3e* and *Ptprc*.
5. LP_L was the only module found to be enriched in differential genes (down expressed in HP) and non-neuronal cell markers (oligodendrocyte precursor cells). The hub genes from LP_L were only found to be enriched in a single gene ontology: *vesicle-mediated transport in synapse* driven by the genes *Prepl* and *Prkaca*.
6. The HP_8/LP_F are modules enriched in genes associated with inhibitory gene clusters Inh_1, Inh_3 and Inh_4, but most notably in clusters Inh_3 and Inh_4 (FDR < 10e-40). Cluster Inh_3 characterized by enrichment in: *Drd2, Adora2, Ppp1r1b, Scn4b, Penk, Rgs9, Calb1* and *Gpr88.* Cluster Inh_4 was characterized by enrichment in a somewhat similar set of genes: *Tac1, Drd1, Pdyn, Ppp1r1b, Scn4b, Oprk1, Rgs9, Strn, Gabrd* and *Reln.* Thus, both modules have a similarity to the expression profile of striatal medium spiny neurons (see e.g. [8]). Both the HP_8 and LP_F modules had no enrichment in differential genes. For the top 25 hub nodes in these modules, the overlap was 72%. Key nodes in this grouping included: *Adcy5, Drd1, Drd2, Gpr88, Pde10a, Ppp1r1b, Rgs9* and *Strn.* The mean kWithin, density, centralization and heterogeneity parameters were similar in these two modules.
7. Inhibitory clusters Inh_4,11 and 12 were *Crh* positive, with cluster 11 being the most enriched; the genes enriched in this cluster are noted above. *Crhr1* positive clusters included inhibitory clusters Inh_88, 10, 13, and 16, while *Crhr2* was expressed in Inh_7 and Inh_9. *Prkcd* CeA neurons have been found important in some ethanol related behaviors (see [29] and references therein). These neurons were detected in a single inhibitory cluster (Inh_2).

**Figure 8.**
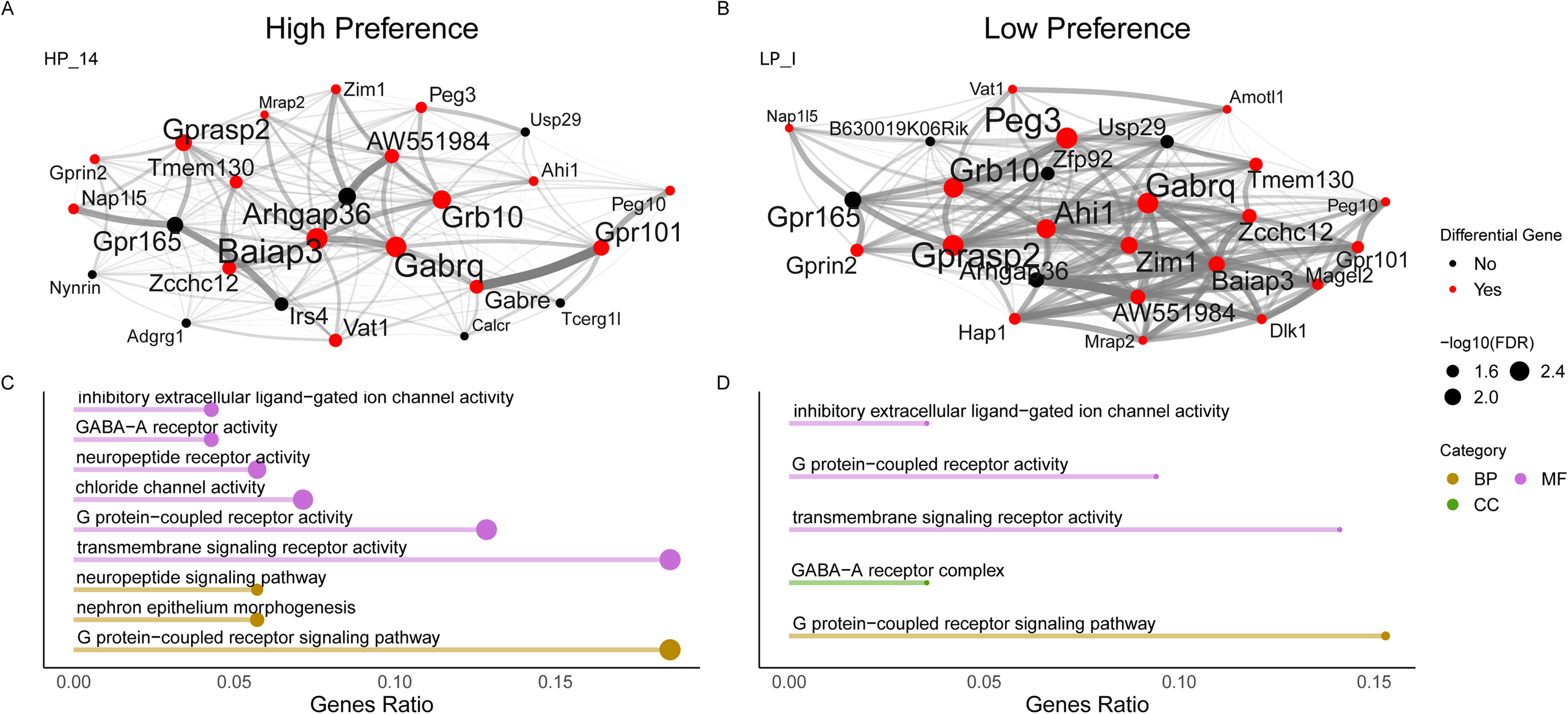
Conserved inhibitory module: A visualization of an inhibitory module conserved across the high preference (HP; left) and low preference (LP; right) networks (Jaccard Similarity = 0.57). The top row (A and B) shows module hub gene connectivity for the top 25 genes in a module (by intra-module connectivity). Lines between hub genes indicate gene-gene connectivity in both opacity and thickness, the size of an individual hub indicates its scaled intra-module connectivity. Hub genes that are differential (DE/DV or DW) are shown in red. Note, a majority of hub genes in each module are differential genes and we can observe a marked drop in module connectivity in the HP module (A) compared to the LP module (B). The bottom row (C and D) shows gene ontology enrichment of the module’s hub genes where the length of the bar indicates the ratio of hub genes in the module to the genes in the GO-term, the size of the dot indicates the significance (-log10(FDR)), and the color indicates to which GO category the enriched term belongs. GO terms are clustered by semantic similarity to reduce redundancy. The HP module (N=11) has more enriched GO terms compared to the LP network (N=5). All GO terms enriched in the LP are also enriched in the HP module (where *GABA-A receptor complex* from the LP terms was clustered with *GABA-A receptor activity* in the HP terms).

**Figure 9.**
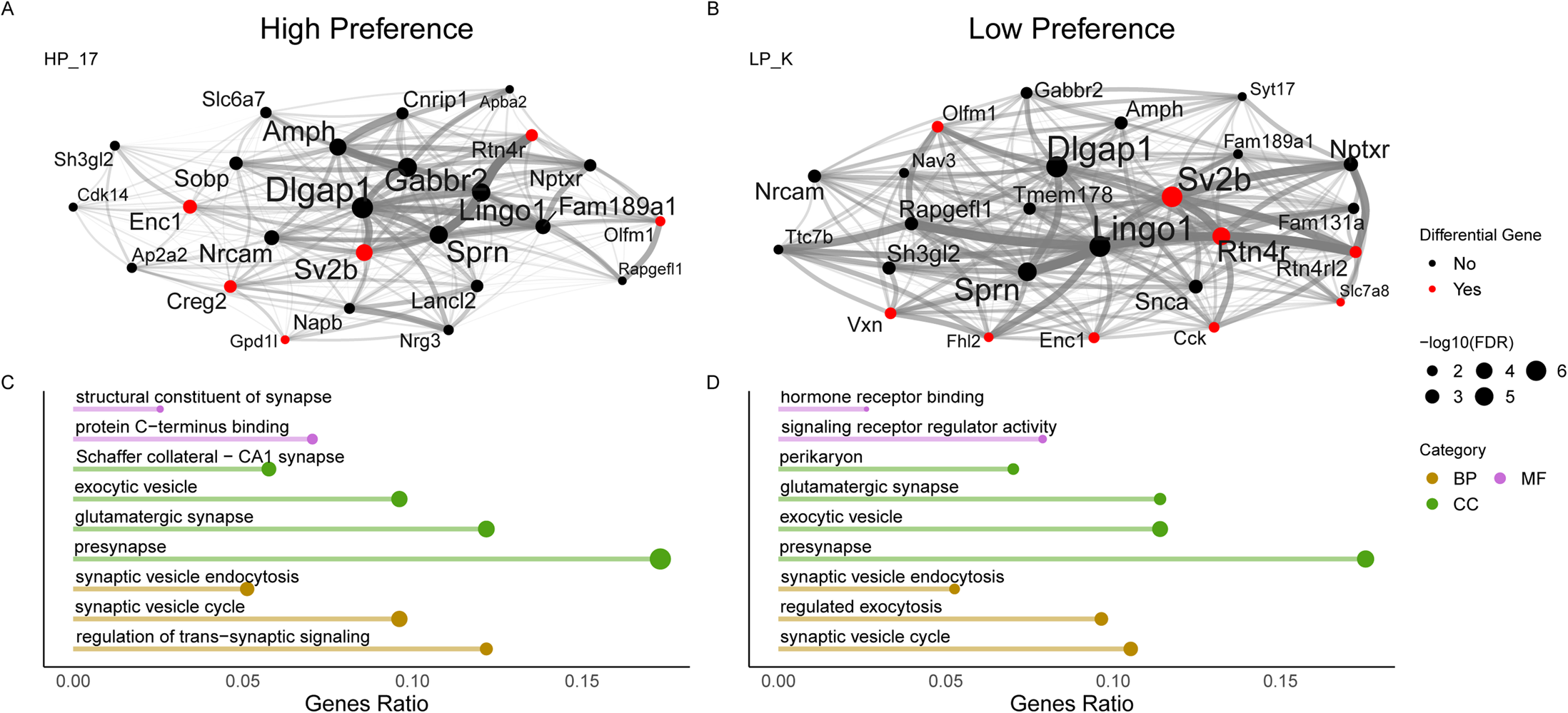
Conserved Excitatory module: A visualization of an excitatory module conserved across the high preference (HP; left) and low preference (LP; right) networks (Jaccard Similarity = 0.47). The top row (A and B) shows module hub gene connectivity for the top 25 genes in a module (by intra-module connectivity). Lines between hub genes indicate gene-gene connectivity in both opacity and thickness, the size of an individual hub indicates its scaled intra-module connectivity. Hub genes that are differential (DE/DV or DW) are shown in red. We can observe a marked drop in module connectivity in the HP module (A) compared to the LP module (B). The bottom row (C and D) shows gene ontology enrichment of the module’s hub genes where the length of the bar indicates the ratio of hub genes in the module to the genes in the GO-term, the size of the dot indicates the significance (-log10(FDR)) and the color indicates to which GO category the enriched term belongs. GO terms are clustered by semantic similarity to reduce redundancy. The HP module (N=39) has more enriched GO terms compared to the LP network (N=21).

## 3 Discussion

Selection for ethanol preference or intake phenotypes has been an integral component of alcohol research for > 70 years. Successful selection studies are found across species (mice and rats), across a variety of genetic backgrounds (outbred populations, heterogeneous stock and F_2_ populations) and many results (behavioral and genomic) have been confirmed in large panels of inbred strains and recombinant inbred panels (see [10] for details). When the selection genetic background is relatively simple e.g. B6 x D2, the results have been remarkably similar (see e.g. [14, 30]) e.g. the same quantitative trait loci (QTL) are reliably detected. However, it is reasonable to ask if one markedly increases the genetic diversity of the founder population (e.g. the HS-CC), how are selection outcomes affected. The current study addresses the outcome issues from two perspectives: 1) Are the HS-CC selected lines behaviorally similar to selected lines derived from simpler founders and 2) Are the genomic mechanisms associated with ethanol preference selection similar. We focus first on the behavioral data.

Mice were bidirectionally selectively bred for high vs. low ethanol preference from a population of HS-CC mice, replicating a prior selection [7]. HP mice from the last selection generation (S4) exhibited a 9.3-fold greater preference for 10% ethanol and a 7.6-fold greater ethanol intake from the 10% ethanol concentration, compared to LP mice. Prior selective breeding projects in mice have used intake as the selection trait (g/kg; [31, 32][31], Phillips et al. [32]) or in one case (with replication), blood ethanol concentration [33, 34], rather than preference. However, Belknap et al. [35] did successfully breed for preference from an F2 cross of two well-characterized, high and low ethanol preferring inbred strains, B6 and DBA/2J, respectively. The lines from the Belknap et al. [35] selection differed 2.0-fold in preference by S4. Thus, a much larger difference arose in the current selection from a highly genetically diverse panel of 8 inbred strains. We cannot be certain of whether this large difference between the two selections would be seen if the lines were bred side-by-side, but it is possible that there are additional genes in the HS-CC for both high and low preference accounting for the larger 9.3-fold difference. The D2 strain is not represented in the HS-CC, but the B6 strain is and furthermore, the PWK strain in the HS-CC has been identified as a high ethanol preference/intake strain [16], with genes that may impact risk for ethanol consumption that are different from those in the B6 strain [17]. Thus, both shared and some different mechanisms likely lead to differences in ethanol intake across populations with different genetic constitutions.

Based on individual values across all HP and LP mice, ethanol intake was highly phenotypically correlated with preference in our selection study (r=0.94). A previous study in a panel of recombinant inbred mouse strains derived from the B6 and DBA/2J strains, found a genetic correlation, based on strain means, of r=0.97 [36]. Therefore, it is likely that these two traits share common genetic influence. However, neither ethanol preference nor intake were strongly correlated with total volume consumed. Overall, S4 LP mice consumed more total fluid than HP mice during testing for selective breeding, indicating that higher ethanol consumption is not due to a general tendency to consume more fluid.

Data for the consumption of fluids with sweet and bitter tastes also did not support a consistent difference between the lines in overall fluid consumption. For the bitter tastant quinine, there was no difference between HP and LP mice in preference nor intake, indicating that differential bitter taste sensitivity does not explain differential ethanol intake. There were no differences between the lines in total volume consumed in the quinine study. HP mice did exhibit greater preference and intake for some concentrations of the sweet solutions, a finding consistent with a body of research showing greater avidity for sweets in correspondence with higher ethanol intake in rodents and in humans that exhibit higher alcohol use or have higher risk for use [37–47]. However, data do not support this relationship across all studies [36, 48–50]. During saccharin and sucrose access, the HP mice consumed more total volume than LP mice, likely as a consequence of their overall greater avidity for the saccharin and sucrose solutions.

Additional traits examined in the HP and LP mice included blood ethanol elimination rate, and sensitivity to the effects of ethanol on locomotor activity and body temperature. Although there were some time-specific sex differences in blood ethanol levels, elimination rate was comparable for the HP and LP mice. Thus, differential rate of ethanol clearance does not provide an explanation for differences in ethanol preference/intake. Likewise, there was no difference in magnitude of hypothermic response between the lines. Ethanol-induced locomotor stimulation in mice has been used as a surrogate for euphoric or activating effects of ethanol found in humans. Whether this initial response to ethanol might predict risk for excessive use or might be experienced at a higher or lower magnitude in risky vs. light drinkers has been considered [51, 52], with heightened sensitivity to stimulant effects found with higher risk for drinking in some studies [53, 54], though not supported in a meta-analysis, in which a moderate (though not significantly different) effect in the opposite direction was found [52]. In our study, LP mice exhibited greater sensitivity to the locomotor stimulant effects of ethanol, compared to HP mice. This finding is consistent with a prior short-term selective breeding project for ethanol intake (g/kg) from the F2 cross of B6 and D2 mice in which the low ethanol consuming line was also more stimulated by ethanol than the high consuming line [32]. These data suggest that some common genes determine ethanol intake and the stimulant response to ethanol. However, differences in sensitivity to the acute stimulant effects of ethanol have not consistently been found in mouse lines bred for differences in ethanol intake [55, 56].

Overall, the behavioral data suggest that the selection for ethanol preference from HS-CC mice is generally similar to that described for selections from less genetically diverse populations. Of course, we recognize that we only sampled a small aspect of potential behaviors. Thus, it seems reasonable that there are behaviors specific to a foundation population, which have not been addressed.

The genetic and genomic features associated with ethanol preference have been recently reviewed [10]. The data suggest that there is overlap across studies. For example, in studies involving crosses between the B6 and DBA/2J strains, the same quantitative trait loci have been reliably detected for 30 years (see e.g. [14, 30]). Colville et al. [7, 8] found that HS-CC preference selection was associated with a gene cluster that had *Dlg2* as a hub node. *Dlg2* encodes PSD-93, which interacts with a number of genes and gene products associated with glutamate receptor activity including *Dlg4, Syngap1, Neto, Grin1, Grin2b, Dlgap1,* and *Dlg3* (see Figure 1 in [8]). Overall, for the LP animals, the data pointed to a strengthening of glutamate connectivity in the CeA and related reward circuitry regions. Similarly, Bell et al. [57] have noted that glutamate connectivity is key to understanding preference in P and NP rats (derived from an outbred Wistar population). With these observations in mind, the current study was undertaken in part to ask how replicable the Colville et al. [7, 8] findings are. To our knowledge the current study is one of the largest to examine the relationships between selection and gene expression and the first (selection study) to integrate bulk and snRNA data. The bulk RNA sample sizes (50/line/sex) were developed empirically to provide co-expression modules with high preservation (see Methods) and the ability to create independent networks for the HP and LP lines (as opposed to creating a consensus network – see [7]).

As no sex by line interactions for differential genes were identified, contrasts for DE, DV and DW were constructed collapsed on sex to focus on selected line differences. Sex will be considered in future publications. Given the large sample sizes (97 HP; 96 LP), it is not surprising that the number of DEGs (FDR < 0.05) was large (2,996). However, many of the significant changes were quite small (1,813 were < 5%). Although the expression of GABAergic, glutamatergic and other receptor and receptor-related genes was increased and decreased in the HP line compared to the LP line (see Results), there was no enrichment in genes for a particular receptor class. The GO analysis of the DEGs (Supplemental Table S10) revealed a significant ontology (FDR < 0.02) in receptor ligand activity but to our knowledge none of the genes (N=21) in this category have been implicated in mechanisms of ethanol action.

As noted in Supplemental Table S6, some of the network modules are enriched (FDR < 0.05) in DEGs while others are not. Here we remind the reader that the DEGs providing an ontology for the network modules are a smaller number than the total number of DEGs. As noted in the Methods, the WGCNA occurs in two steps. The first step utilizes all 14,672 of the genes detected as present. This step allows one to rank order the genes in terms of network connectivity. The genes contributing the bottom 10% of network connectivity are culled leaving 10,716 genes. The culled genes are those with a low variance. This culling step reduces the number of DEGs associated with the network to 1,666.

Replication is a key step in scientific rigor, and we took this step by repeating our selection of LP and HP lines from the HS-CC founder. This provides the opportunity to ask whether the DEGs associated with the specific network modules we identified provide significant new information? The answer to this question was mixed but overall, there was a general consistency with previous findings. Of the HP modules, only the DEGs up-regulated in HP_15 (FDR < 10^-24^) exhibited significant gene ontologies. Significant results (FDR < 0.05) were detected for the related ontologies of dendrite, synapse part, synapse, neuron part, neuron cell body, cell body and intrinsic component of synaptic membrane (Supplemental Ontology Tables). Similar results were obtained for DE up-regulated genes in module LP_B (Supplemental Ontology Tables). This LP module is the most similar to HP_15, although the Jaccard similarity was low (0.20).

Differential variability was a key metric in the Colville et al. [7, 8] analyses. Here we note that all of the significant DV genes were retained in the parsed WGCNA networks (see above). DV was lower by a 10:1 ratio in the HP network. However, we recognize that the loss of connectivity will to some degree reflect that technical artifacts such as batch effects mask the residual gene connectivity. Supplemental Table S10 indicates that the DV genes have a rich ontology and include positive regulation of synaptic transmission, modulation of chemical synaptic transmission, transmembrane receptor protein kinase activity and synaptic transmission (cholinergic). Genes associated with the positive regulation of synaptic transmission (FDR < 10^-5^) included ***Tacr1, Adora1, Ntrk2, Arc, Adrb1, Htr2c*** and ***Chrnb2***. Notably, and in contrast to Colville et al. [7, 8], *Dlg2* was not a DV gene. It is important to note as we have previously found (see [8]), that the relationship between variance and network connectivity is an inverted “U” function, assuming the changes in variance have biological significance. Thus, as variance increases, connectivity will increase up to the point where the change in variance becomes chaotic and loses biological relevance.

Differential wiring assesses large changes in the edges among individual genes and defines a unique subset of gene network-related changes. Of the 407 DW genes, 334 had 5 or fewer edge changes. In the HP line, 283 (70%) of the DW genes showed a decrease in wiring. Significant GO terms associated with the DW genes include cytosolic ribosome, cytosolic small ribosomal subunit and neuropeptide receptor activity. Genes in the latter category include ***Prlhr, Tacr1, Qrfpr,*** *Sstr5, **Galr1, Oprm1,*** and ***Prokr2*** (see Supplemental Table S10). All of these genes with the exception of *Sstr5*, exhibited decreased wiring in the HP line. Of note, *Oprm1* is predicted to have a prominent role in AUD [58–60] and ethanol preference [61, 62]. *Oprm1* edges affected included those with *Arc, Cdk2ap1, Ephb6, Qrfpr, Trib1,* and *Unc5d.* However, Kaoru et al. [63] found that *Oprm1* is associated with the intercalated cells, which surround the periphery of the BLA [62, 63]. The BLA does project to the CeA [26], suggesting the possibility that the gene may be expressed in projection neurons terminating in the CeA. It is also possible that our *Oprm1* findings are an artifact of the procedure used to punch the CeA, but may highlight the relevance of the BLA to ethanol consumption (e.g. [64]).

As noted above, the sample sizes (∼100/line) allowed the formation of separate networks for the HP and LP lines as opposed to the consensus network approach used previously [7, 8]. A comparison of the HP and LP networks revealed both significant similarities and differences. The differences between the networks e.g. only 6 pairs of the HP and LP modules have a significant Jaccard similarity could be due to multiple causes. Genetic drift in the networks may have differed. However, we note that short-term selective breeding is designed to minimize the drift effects. In addition, breeding a large number of families/line (N=24) should minimize random drift effects. Nonetheless, we recognize that some of the network differences may not be selection related. A current focus of the analysis assumes that differences in DE, DV, or DW are due to selective breeding, and represent significant contributions to phenotypic differences between HP and LP animals. However, we note that exon-level expression or alternative splicing may also be driving the selection phenotypes. Given the large amount of data generated and the extensive analyses already performed, we plan to explore exon usage and splicing variation in future work.

We focused our analyses on the three pairs of synaptic modules that are preserved in both lines, and also are significantly enriched with genes associated with the effects of selection: HP_14/LP_I; HP_17/LP_K; HP_8/LP_F. Of the three pairs, only one (HP_14/LP_I) showed a significant difference in network connectivity between the LP and HP lines (∼80% lower in the HP line). This difference was detected when all module members were included in the analysis or when just the hub nodes were included (see Supplemental Table S6). A decrease in module connectivity is here interpreted to indicate that the module is working less efficiently within, will be less efficiently connected to other modules, and will be less robust to perturbations. This sort of modular disruption is thought to be in part responsible for a variety of disease states including neurodegenerative and neuropsychiatric disorders, cancer and metabolic disorders (see e.g. [65]). However, despite the loss of connectivity in the HP line, the essential elements of the module structure remained intact. The genes in the HP and LP modules are essentially identical as are the hub nodes. Module ontologies are also similar with a notable exception that in the LP module there was a significant enrichment in genes associated with inhibitory extracellular ligand-gated ion channel activity (FDR < 3 x 10^-5^) not found in the HP line (Supplemental Ontology Tables). We included looking for B6/PWK “signature genes” in the analysis of WGCNA modules, but note this approach assumes that DE genes in the HP/LP animals equate with those from comparisons of two high-drinking HS-CC progenitor strains (B6 and PWK) versus the other six progenitors. It is possible that epistatic interactions could differ substantially between the HP/LP lines and the original founders, potentially leading to expression and preference differences not observed in the inbred lines.

To understand the likely functional consequences of the decrease in modular connectivity, snRNA-Seq data were used to determine which neuronal clusters were enriched in the HP_14 and LP_I modules. The data pointed to inhibitory clusters Inh_6 and Inh_18; note that based on cell counts, cluster Inh_6 was more than 10 times larger than cluster Inh_18. As shown in the UMAP (Figure 3A), cluster Inh_18 appears to be a daughter cell type to cluster Inh_6. Both clusters are enriched in genes associated with *Isl1* inhibitory clusters. O’Leary et al. [23] found that the *Isl1* expressing neurons were a previously unresolved population of CeA neurons, largely associated with the medial aspect of the CeA. Immunohistochemistry and long-range projection mapping were used to show that these neurons project prominently to the substantia nigra, the parabrachial nucleus and the periaqueductal gray; the *Isl1* cells also project non-specifically to other brain regions, including the hypothalamus (see Figure 8 in [23]). The *Isl1* clusters (Inh_6 and Inh_18) appear to be differentiated from other commonly used cluster markers such as *Drd2* (clusters 3, 8 and 16), *Drd1* (clusters Inh_1 and Inh_4), *Crh* (clusters Inh_4, Inh_11 and Inh_12), *Nts* (Inh_11), *Prkcd* (cluster Inh_5) and *Nr2f2* (clusters Inh_6, Inh_8, Inh_13 and Inh_14). Hochgerner et al. [22] observed two clusters of *Isl1* neurons, one enriched in *Tac1* and the other enriched in *Aldoc* (adolase, fructose-bisphosphate C); we confirm enrichment in these clusters (Inh_18 and Inh_6). We also confirm that the *Prkcd* cluster (linked to ethanol withdrawal) is highly enriched in the expression of *Sox5* and *Ezr*, but we find less specifically enriched in *Fgfr2* (see [29]) or *Oprk1, Adora2* and *Nts* (see [22]). The integration of modules 14 and I with the expression of inhibitory clusters Inh_6 and Inh_18 is perhaps most clearly seen in shared gene membership. For the HP_14 module, 43 genes identified as markers (top 10% by rank) for inhibitory cluster Inh_6 have membership, and 56 genes identified as markers for inhibitory cluster Inh_18 have membership (Supplemental Table S15). Of the genes with shared membership for Inh_6, 23 (46%) were hub nodes and while 26 (38%) were hub nodes for Inh_18. The parallel data (Supplemental Table S16) for the LP_I module are 47 genes (20 hub, 43%) with shared membership for cluster Inh_6 and 64 genes (28 hubs, 44%) shared with cluster Inh_18. Given the extent of the overlap between the modules and gene clusters, we assume the HP gene clusters will be similarly less efficient and robust to external perturbation. The key piece of information that we cannot extract from the current data is whether the effects of ethanol on these inhibitory neurons will also be less effective.

The data described above on the differential genes (expression, variance and wiring) and the network analysis points to substantial effects of selection on the CeA transcriptome and circuitry. However, it was also observed that some network modules were preserved in the HP and LP lines. This preservation may be random, but the alternative view is that the preserved modules have a high functional value. Of the six preserved module pairs, a significant effect of selection (down in the HP for DE, DV and DW) was found on only one pair enriched in inhibitory cell markers – HP_14/LP_I, in which HP connectivity was also markedly diminished. These modules were enriched in genes associated with the *Isl1* neurons that project from the central medial nucleus of the central amygdala. The preferred *Isl1* projection targets (substantia nigra, parabrachial nucleus and the periaqueductal gray) are all regions that have been linked in some way with ethanol consumption [66–68]. It is of interest to note that both clusters Inh_6 and Inh_18 are enriched in tachykinin associated genes, including *Tac1*, *Tac1r* and *Tac3r*. Tachykinins have been implicated in genetic risk for ethanol use [69]. *Tac1* encodes the neurokinin peptide Substance P and other peptides in the tachykinin family of hormones; Substance P binds to the neurokinin 1 receptor expressed by *Tacr1*; neurokinin B binds to the receptor expressed by *Tacr3*. There is some precedent for an association of *Tacr1* expression in the CeA with risk for high ethanol intake. Thus, *Tacr1* expression was elevated in the amygdala of alcohol-preferring P rats compared to progenitor Wistar rats and this was accompanied by greater neurokinin 1 receptor binding in the CeA of P rats [70]. Upregulation of *Tacr1* in the CeA resulted in escalation of ethanol self-administration in P rats [71]. In B6 mice, known to have a high propensity for ethanol preference and intake compared to other strains of mice, and in rat lines bred for high ethanol intake, inhibition of neurokinin 1 receptor signaling via reduced *Tac1r* expression, pharmacological antagonism or genetic deletion of *Tacr1* reduced ethanol consumption [70, 72–75]. However, in one report, lower *Tac1* mRNA levels were found in the CeA of P compared to NP rats and Substance P reduced ethanol responding when infused into the CeA of P rats [76]. Whether tachykinins are a source of differential risk for ethanol preference and intake between the HP and LP lines is something that remains to be explored.

In sum, we find that five generations of bidirectional short-term selective breeding for ethanol preference from HS-CC reproducibly produces lines of mice that prefer and consume ethanol significantly differently from each other. The effects of selection in naïve mice from generation S5 on the transcriptional risk vs. protection signature for ethanol drinking in the CeA are profound, with bulk RNA sequencing identifying DE in ∼18% of genes and a marked decrease in gene expression variability and connectivity in ∼3% of genes. While no differences in cell-type specific expression nor in the proportion of cell-types between the HP and LP lines were identified using snRNA-Seq analysis, marker genes were identified that provide high specificity for distinguishing inhibitory and excitatory neuronal sub-types from one another. By leveraging high-powered bulk RNA-Seq data to construct robust gene co-expression networks, we were able to identify networks of genes that were highly associated with the effects of selection and preserved within both the HP and LP mice, and then relate these to specific neuronal sub-types. These findings, which have overlapping lines of evidence across data from different modalities, provide a powerful tool for further hypothesis generation to identify potential interventional and mechanistic studies.

## 4 Methods

### 4.1 Animals

All experiments were performed in accordance with the National Institutes of Health Guidelines for the Care and Use of Laboratory Animals and were approved by the Institutional Animal Care and Use Committee of the VA Portland Health Care System. Mice were housed in shoebox cages with Bed-O-Cob bedding and maintained at 21 ± 1°C on a 12-hour light:dark cycle, with lights on at 0600 h. Behavioral testing occurred between 0800 and 1400 h. Mice had ad libitum access to tap water and standard rodent chow (Purina 5001 or 5LOD Pico Lab Rodent Diet; Animal Specialties, Woodburn, OR, USA), except when noted below. Mice were weaned at 21 + 2 days and group-housed with same-sex littermates in groups of two to four mice.

### 4.2 Drugs and Other Compounds

Ethanol (200 proof) was purchased from Decon Laboratories, Inc. (King of Prussia, PA, USA) and mixed with tap water for consumption. For IP injections ethanol (200 proof) was mixed with sterile saline (0.9% NaCl; Baxter, Deerfield, IL, USA).

### 4.3 Selective Breeding for High and Low Ethanol Preference

A population of 187 HS-CC mice (94 female and 93 male) served as the founding population for the second replicate HP and LP selected lines; data for the initial set of lines have been reported [7, 8]. The HS-CC stock was established within the VA Portland Health Care System veterinary medical unit by pseudo-random interbreeding of 8 inbred strains (129Sv/Im, A/J, C57BL/6J, Cast/Ei, NOD/Lt, NZO/HILt, PWK/Ph, WSB/EiJ), chosen to fully represent Mus musculus genetic diversity. The breeding population comprised 48 families bred using a rotational breeding scheme, but is no longer maintained. The founders for the HP and LP lines were from generation 41.

The selective breeding trait was preference for 10% v/v ethanol in water offered vs. water in a continuous access, two-bottle choice procedure. At 7-8 weeks of age, mice were habituated for one week to individual housing. After this period, two 25-ml water-filled graduated cylinders fitted with rubber stoppers and sipper tubes were placed on each cage for 4 days to familiarize the mice with drinking from sipper tubes; amount consumed from each tube was measured daily (± 0.2 ml). On days 5-8, a water tube was replaced with a tube containing 5% ethanol in water (v/v) and on days 9-12, water was offered vs. 10% ethanol (v/v). Amount consumed was measured daily. The relative positions of the water and ethanol tubes were switched every two days to control for side preferences. Tubes containing the same solutions were placed on empty cages to measure leakage and evaporation, and average volume lost from these tubes was subtracted from individual drinking values. Dependent measures were g/kg ethanol consumed, based on body weight obtained every 4 days; total volume consumed (ethanol in ml + tap water in ml); and ethanol preference ratio (ml of ethanol consumed/ total fluid consumed in ml).

Average 10% ethanol preference ratio on the two days after a position switch (days 10 and 12) was used as the selection phenotype, consistent with replicate 1 [7]. The HP line was established by interbreeding the 20-24 pairs of males and females with the largest preference ratio. The LP line was established by interbreeding the 20-24 pairs of males and females with the smallest preference ratio. In subsequent generations, offspring were produced by 20-24 successful breeding pairs per line and mice were isolate housed at 8 weeks of age and then tested as described for the HS-CC mice; 183-257 total offspring were tested each generation. In addition to ethanol preference, g/kg ethanol intake and ml/kg total volume consumed were measured and analyzed.

#### 4.3.1 Blood Ethanol Elimination Rate

To examine ethanol elimination rate, mice were moved to a procedure room, weighed, and returned to their home cages for 1 h before study initiation to allow acclimation to the new environment. Each animal was then injected IP with 2 g/kg ethanol (20%, v/v) and 20 µl blood samples were obtained from the lateral tail vein 15, 30, 60, 120 and 180 minutes after administration. To obtain blood samples, mice were placed into cylindrical acrylic plastic restrainers from which the tail protruded, the tail tip was removed with sharp scissors (< 1 mm), and the tail was gently milked, and each sample exuded into a microcapillary tube. The blood was aspirated into glass vials on wet ice, containing 0.0003% propanol solution. After each sample collection, the vial was immediately capped, the cap was crimped to create a tight seal, and the vial was lightly inverted a few times and returned to wet ice. After all samples were collected, they were frozen at −20°C until assayed by gas chromatography, using well-established methods [77, 78]. Selection generation, age, and group size are provided in the figure legend for this study.

#### 4.3.2 Other Behavior Traits

Methods for studies examining voluntary consumption of sweet and bitter tasting solutions and sensitivity to the effects of ethanol on locomotor activity and body temperature are provided in the Supplemental information.

#### 4.3.3 Behavioral Data Analysis

Data were analyzed by factorial ANOVA, with repeated measures when appropriate. Significant interactions involving multiple factors were deconstructed to identify sources of interaction. Two-way interactions were interrogated using simple main effects analysis followed by Newman-Keuls post hoc mean comparisons. All analyses initially included sex as a factor. Follow-up analyses were conducted with data from the sexes combined when sex did not interact with other factors. Effects were considered significant at α ≤ 0.05. Behavioral data analyses were performed using Statistica 13 software (Tibco Software, Inc., Palo Alto, CA, USA).

### 4.4 RNA Sequencing

To investigate the transcriptome preceding exposure to ethanol and thus, contributing to risk for or protection from high ethanol preference and intake, we performed bulk RNA sequencing on tissue collected from the CeA of HP and LP mice (n ∼50/sex/line; N=200). To determine if selection altered the distribution of CeA cell types and provide cell-type ontology information, snRNA sequencing was performed on CeA tissue collected from 16 S5 generation mice (n = 4/line/sex). The methods used for bulk and snRNA-Seq were motivated by current best practices [79–84]. In addition, we took guidance from recent works utilizing snRNA-Seq data from the CeA of B6 mice, including seminal efforts of The BRAIN Initiative Cell Census Network (BICCN) to map the whole mouse brain using snRNA-Seq and spatial transcriptomics [22–25].

#### 4.4.1 Tissue Processing

##### Dissection

Brain tissue was obtained from experimentally naïve, non-ethanol exposed, S5 HP and LP mice. Mice were 60-70 days of age at the time of tissue harvesting (mean ± SEM = 63 ± 2 days). Mice group-housed by sex (2-4 per cage) were moved to a procedure room at 1000 h and allowed to acclimate for one hour; brains were obtained at 1100 - 1300 h. Mice were euthanized via cervical dislocation immediately followed by decapitation. Brains were removed using RNase-free tools, immediately frozen on dry ice, placed in RNase/DNase free polypropylene tubes and stored at −80°C until dissected. Frozen brains were dissected using anatomical landmarks from Franklin and Paxinos [85]. The CeA was obtained bilaterally, using RNase free tools, an aluminum dissection stage, and micropunches fashioned from blunt-cut 27 gauge needles. The frozen brain was placed ventral side up on the dissection stage and cut using a razor blade just anterior to the posterior boundary of the hypothalamus at bregma −0.82 mm. A second cut posterior to the first was then made at bregma −1.46 mm to obtain a 0.6 mm slice. The CeA was removed from this slice using the forked end of the external capsule, the lateral globus pallidus, and the internal capsule as landmarks. The bilateral CeA punches for each animal were placed in RNase and DNase free microcentrifuge tubes and stored at −80°C until RNA was extracted.

##### Bulk Extraction and Sequencing

Frozen tissue samples were transferred to the Gene Profiling Shared Resource at Oregon Health & Science University for total RNA isolation, using QIAsymphony RNA extraction methods for fatty tissue (QIAsymphony RNA Handbook, November 2020; Qiagen, Redwood City, CA, USA). Quality of the RNA was assessed by UV absorption (NanoDrop 8000) and all CeA samples had an average 260/280 ratio of 2.3, indicating high sample purity. RNA Integrity Number scores were obtained using a Bioanalyzer assay and ranged from 9.1 – 10.0, consistent with high quality RNA. Samples were transferred to the Massively Parallel Sequencing Shared Resource (MPSSR) for RNA-Seq data collection. The total number of samples was 200; 47-52/line/sex. The MPSSR completed library formation and sequencing according to Illumina’s specifications. Stranded libraries (polyA+) were prepared using the TruSeq Stranded mRNA kit (Illumina, San Diego CA, USA) and were multiplexed in five batches of 40 on an Illumina NovaSeq 6000, yielding approximately 20 million paired-end reads per sample. Batches were balanced for sex and line.

##### Single Nuclei Extraction and Sequencing

snRNA-Seq libraries were generated from 16 S5 generation mice (n=4/line/sex). Flash-frozen CeA was dissociated into individual nuclei following manufacturer’s instructions (protocol CG000366, 10X Genomics). Nuclei were filtered through a 40-70 μm mesh filter and centrifuged in a 29-50% OptiPrep gradient (Sigma-Aldrich) to separate intact nuclei from debris. Cells were inspected in a counting chamber for intact, bright, nongranular cell morphologies, indicating high viability and successful debris removal. ∼ 10,000 single nuclei were captured for each sample in a single channel on the 10X Chromium controller, and snRNA/ATAC-Seq libraries (Chromium Next GEM Single Cell Multiome ATAC + Gene Expression) were generated following the manufacturer’s instructions (10X Genomics). Libraries were sequenced to an average depth of 20,000-25,000 read pairs per nucleus (RNA-Seq and ATAC-Seq, respectively), in a NovaSeq6000 at the MPSSR according to 10X Genomics’ specifications. All samples were processed and sequenced at the same time to avoid batch effects. Before filtering and clustering, we captured a total of 87,570 cells with a median of 4,890 UMI counts per cell and 2,320 genes per cell.

#### 4.4.2 Data Cleaning - snRNA

##### Alignment

Raw sequencing data for both the ATAC and RNA were demultiplexed, aligned to the GRCm39 mouse genome (Ensembl Mouse 111; GRCm39 GCA_000001635.9), and both mRNA molecules and ATAC peaks were counted with the cell ranger arc pipeline (10x Genomics, v 2.0.2). RNA transcripts were assigned to cell and molecule of origin using barcode sequences from the bead primers. The snRNA-seq data are available via NCBI Gene Expression Omnibus (accession number GSE293636).

##### Droplet Filtering

Raw RNA and ATAC libraries for each sample were imported individually in RStudio using R version 4.4.2 as Seurat [86] objects (v5.1.0). 8.4M droplets (∼524k per sample) with nonzero UMI and genes were sequenced. As the primary focus for this manuscript was mRNA, we discarded the ATAC data at this point for the sake of brevity. The ATAC data will be analyzed at a later date, along with additional whole genome sequencing data. Distributions of unique molecular identifiers (UMI; counts) and uniquely mapped genes (features) were examined, and a lower bound of 250 UMI was used to separate empty droplets from nuclei-containing droplets, resulting in 100,174 remaining.

These distributions were observed to be bimodal, showing two populations of droplets with one group containing an average of ∼1,000 distinct genes (∼2,000 UMI) and the other group containing an average of ∼4,000 distinct genes (∼10,000 UMI). Samples were examined individually to determine cutoffs to eliminate high and low tails for counts and features for the raw, unnormalized data. A full annotation of snRNA-Seq samples is available in Supplementary Table S17, but the mean low feature cut off was ∼600 unique genes (∼1,200 counts), and the mean high feature cut off was ∼6,700 unique genes (∼28,000 counts).

Next, any droplets meeting the following criteria were dropped: (1) more than 0.5% mitochondrial RNA; (2) more than 10% of total counts belonging to a single gene; (3) low complexity (ratio of features to counts of greater than 0.85). After filtering, 69,318 droplets (hereafter referred to as cells) remained, with ∼47k having ∼4,000 distinct genes, and ∼22k having ∼1,000 distinct genes.

##### Gene Filtering

After filtering for low-quality and empty droplets, a total of 33,589 genes had non-zero counts. As all downstream analysis will focus on a relatively low number (thousands) of highly variable genes, we wish to filter out low expressing genes for the sake of computational and memory efficiency. We drop any genes which have a count of 2 in less than 10 individual cells from further analysis. We additionally follow Hochgerner et al. [22] and drop the following immediate early genes and sex-genes (*Btg2*, *Jun*, *Egr4*, *Fosb*, *Junb*, *Gadd45g*, *Fos*, *Arc*, *Nr4a1*, *Npas4*, *Coq10b*, *Tns1*, *Per2*, *Ptgs2*, *Rnd3*, *Tnfaip6*, *Srxn1*, *Tiparp*, *Ccnl1*, *Mcl1*, *Dnajb5*, *Nr4a3*, *Fosl2*, *Nptx2*, *Rasl11a*, *Mest*, *Sertad1*, *Egr2*, *Midn*, *Gadd45b*, *Dusp6*, *Irs2*, *Plat*, *Ier2*, *Rrad*, *Tpbg*, *Csrnp1*, *Peli1*, *Per1*, *Kdm6b*, *Inhba*, *Plk2*, *Ifrd1*, *Baz1a*, *Trib1*, *Pim3*, *Lrrk2*, *Dusp1*, *Cdkn1a*, *Pim1*, *Sik1*, *Frat2*, *Dusp*5, *Xist*, *Tsix*, *Eif2s3y*, *Ddx3y*, *Uty* and *Kdm5d*) from further consideration, leaving 14,533 for downstream analysis.

#### 4.4.3 Data Analysis - snRNA

In order to maximize our ability to robustly identify cell sub-types and markers, we employed an iterative clustering and cell type-annotation process following the logic of other recent works exploring snRNA-Seq in the CeA [22–24].

##### Normalizing and Integration

Samples were individually normalized using the *SCT*ransform function in Seurat, which uses analytical Pearson residuals for transformation and is known to perform particularly well for clustering and marker gene selection in the presence of rare cell populations [80, 87–89]. All samples were then merged into a single *Seurat* object, and count corrections were computed to normalize depth across all samples (in contrast to the commonly used *cellranger aggregate* pipeline, which randomly downsamples to normalize sequencing depth across libraries). No differences were observed in cluster composition in regard to genotype, sample, sex, or sequencing lane, and so no integration or batch-correction was performed.

##### Clustering

For each stage of the clustering the following steps were taken. First, principal components were calculated, then used to generate a UMAP (for visualization), to calculate k-nearest neighbors and construct a shared nearest neighbor graph. Next, the Leiden algorithm [90] is then used to identify clusters across 12 distinct resolutions, and a final resolution is chosen via visual inspection of the UMAP as well as dendrograms (constructed using Euclidean distance of normalized gene expression). In general, a resolution which included spatially isolated clusters and was the first resolution to split a major cluster into parts was preferred. In all cases a resolution between 0.25 and 0.8 was used. Markers are then identified using the *Presto* [91] package for Wilcoxon differential expression, where significant markers (FDR < 0.05) were ranked (equally weighted) by their: (1) log-fold change, (2) AUROC metric and (3) relative abundance (log-fold change of the precent cells expressing the gene). Fischer and Gillis [92] found using multiple datasets from the BICCN that AUROC and log-fold change could be used for rapid identification of ideal markers and that 50-200 was ideal for enrichment testing. Here we also included relative abundance as an additional module characterizing and attempt to use between 50-100 markers for enrichment testing. Markers were used to manually annotate clusters, identify and remove doublet clusters and low-quality/noisy clusters. Clusters are always ordered according to their relative total cell count (e.g. Inh_1 has more cell than Inh_2).

Clusters are additionally ordered using unsupervised hierarchical clustering on the Euclidean distance of the highly variant genes to aid visualization and to allow for the identification of clusters which have little differences in their expression patterns and may be combined. This hierarchical clustering also allowed for identification of markers between bifurcations of the associated dendrogram; a positive marker in this context means expression in at least 50% of cells on that side, and a log-fold change of 2 (400% expression), while a negative marker means expression in lower than 5% of cells and a log-fold change of −2 (25% expression). In general, a positive marker on one side is a negative marker on the other, but the opposite is not always true. A positive marker is one which is highly abundant and has higher expression on one side of the tree-cut, while a negative marker is one which is both rarely and lowly expressed on one side of the tree-cut.

##### Coarse Clustering

We first conducted a course clustering as above using all 69,318 cells, and identified clusters using a majority vote of known markers in their aggregate expression as either inhibitory neurons (expressing *Gad1 or Gad2*) or excitatory neurons (expressing *Slc17a7 (*Vglu1) or *Slc17a6 (Vglu2)*). All remaining clusters were considered non-neuronal. At this point two samples were excluded from further analysis, one for having abnormal high proportion of non-neuronal cells (more than 50%) and another for having twice as many excitatory neurons as inhibitory neurons, the compositional makeup of all samples at this stage of clustering is shown in Supplementary Figure S10. Dropping these two samples resulted in a final count of 53,839 cells.

Next, markers were identified which distinguish the inhibitory neurons from all other clusters, the excitatory neurons from all other clusters, and the non-neuronal clusters from the neuronal clusters. Manual annotation and literature review [22–24] yielded the following markers for major cell types: Inhibitory (*Gad1, Gad2, Dlx6os1, Slc32a1*); Excitatory (*Slc17a7, Slc17a6, Nrn1*); Astrocyte (*Gja1, S1pr1, Fgfr3, Gli2, Ntsr2*); Microglia (*Cx3cr1, Ikzf1, C1qa, C1qb, C1qc*); Oligodendrocyte (*Mal, Mog, Mag, Prr5l, Plp1*); Oligodendrocyte Precursor Cells (*Pdgfra, Cspg4, Olig1, Olig2, C1ql1*); Immature Neurons (*Tmem72, Oxt2os1, Mecom, Oxt2, Folr1*); Endothelial (*Vtn, Cldn5, Pdgfrb, Flt1, Acta2, Ly6c1*). These were used to classify, clusters in aggregate, and cells, individually, to identify cells to include in finer clustering as above via majority vote. Potential doublet clusters and cells were identified following the methodology of Hochgerner et al. [22], where the expression of the identified cell type markers is compared to all other types, and if the worst ratio is less than 2 than it is considered to be a potential doublet. At the cell level this is likely to include many false positives, as cells where the key markers were excluded from amplification by random chance will also be identified. However, Yao et al. [25] noted that cells which were missing key markers correlated strongly with other low-quality cell metrics, and we will remove these cells to maximize the quality of cells used in fine clustering. In total, 7,620 (∼14%) cells were identified as potentially low quality in this manner and excluded.

Note, the bimodal distribution of UMI and genes noted in Section 4.4.2 appeared to correspond roughly to neuronal (which had mean counts of ∼12,000 and ∼4,000 genes) and non-neuronal (mean counts of ∼2,000 and ∼1,000 genes) cell populations.

##### Fine Clustering

Cells identified as either inhibitory neurons, excitatory neurons or non-neuronal were clustered separately as above to identify sub-populations and to maximize power and specificity when identifying markers which distinguish each sub-type from its main class. For example, we are most interested in what most distinguishes a given inhibitory neuronal subtype from other inhibitory neurons, rather than all other cell types in the CeA. As before, we drop any genes which have a count of 2 in less than 10 individual cells from further analysis. We additionally drop any cluster which has fewer than an average of 10 cells per sample. A count of cells identified in each cluster after fine clustering (of inhibitory, excitatory and non-neuronal populations) is available for each animal in Supplementary Table S18.

##### Compositional Analysis

In order to determine if the proportional of cell types differs between the high- and low-preferences line a linear model including line and sex was run on centered log ratio transforms of cell-type compositions. Results were FDR corrected across the number of cell-types tested. Clusters with lower than an average of 30-cells per cell-type were excluded.

##### Differential Expression

DEGs were identified using a consensus method proposed by Salem et al. [93] in order to minimize the impacts of *pseudoreplication* [80]. For a given cluster, DEGs were identified if detected (FDR < 0.05) in three of four distinct pipelines. Two *pseudobulk* methods (generated via the *Seurat AggregateExpression* function) were used which employ distinct statistical approaches (*edgeR* [94], and *Deseq2* [95]). Two single-cell methods were used which test the two-part hypothesis of a difference in either magnitude of expression or proportion of expression were also used (MAST two-part hurdle GLMM [96, 97], and *DEsingle* [98]). For single-cell methods animals with fewer than 10 cells in a cluster were excluded, and any gene which had fewer than 10 cells in a line by sex group (e.g. male HP mice) was excluded. Default *edgeR* filtering as per the user guide (for genes, *filterByExpr,* and samples, library size greater than 50,000) was used for both pseudobulk methods. Sex was explicitly included in the model wherever possible (the exception being DEsingle). To reduce computational time, pseudobulk methods were evaluated first and only genes that were significant DE in at least one pseudobulk method were tested in the single-cell methods. P-values in the single-cell methods were then FDR corrected to the number of genes used in the pseudobulk methods. For significantly DE genes, edgeR calculated fold change was reported. DEGs were calculated independently for each cluster, and FDR was applied independently for each cluster and method.

#### 4.4.4 Data Cleaning – Bulk RNA

##### Alignment

FastQC was used for initial quality checks on the raw sequence data and no issues were observed. STAR aligner was used to align the sequenced data to the GRCm39 version of the mouse genome (Ensembl Mouse 105; GRCm39 GCA_000001635.9), allowing for a maximum of three mismatches per 100 base-pairs read. On average, 90% of reads were uniquely mapped. STAR aligner was also used to obtain read counts at the gene level. Raw gene expression counts for 55,414 aligned genes and gene-like features were imported and analyzed in RStudio using R version 4.1.2.

##### Gene filtering

Prior to additional analysis 39,216 genes were excluded (39,115 for having mean counts-per-million below 0.5 across samples; seven for having a count above 120,000 in a single sample; 94 for being non-chromosomal). The annotation for the remaining 16,198 genes (14,287 protein coding, ∼88%) is provided in Supplemental Table S5A, the annotation for the 39,216 dropped genes (7,597 protein coding, ∼19%) is provided in Supplemental Table S5B. Ensembl 105 (released Aug 2021) was used for annotation, matching the version used for aligning, and was retrieved on July 24^th^, 2023 using the *biomaRt* R package. The bulkRNA expression data are available via NCBI Gene Expression Omnibus (accession number GSE288772)

##### Sample filtering

A total of seven samples were dropped from further analysis; three samples for being potentially mis-gendered (as determined by *Xist, Kdm5d, Eif2s3y, Uty,* and *Ddx3y* counts); one sample for being potentially mis-aligned (as determined by principal component analysis); and three samples for having an inter-sample-correlation three standard deviations away from the median.

Categorical information for the dropped samples is provided in Supplemental Table S4A. The dropped samples were evenly distributed among sex and line (F = 4, M = 3, HP = 4, LP = 3). Categorical information for the 193 samples (49 F HP, 49 M HP, 46 F LP, and 49 M LP) included in analysis may be found in Supplemental Table S4B.

##### Filtering limitations

One additional feature of the data warrants discussion here. Nine mice had zero counts in 1% or more of the 16,198 genes included in analysis. There were no detectible trends in which genes these mice had zeros and ∼5% of genes had three or more mice with zero counts. This was originally considered to be a technical artefact, but examination indicates eight (∼90%) of these nine mice were in the low preference line. Since it is unclear if these zeros are biologically important, we included them in the analysis out of an abundance of caution. Neither the mice, nor the genes, have been removed and no zeros were replaced with imputed values. The distribution of zero counts for genes and mice is given in Supplemental Figures S11 and S12.

The categorical information for the nine mice is provided in Supplemental Table S4C.

#### 4.4.5 Data Analysis – Bulk RNA

##### Exploratory Data Analysis

PCA was performed using the *prcomp* function of R’s base stats package on counts normalized using the “trimmed mean of M-values” method as part of the *limma+voom* pipeline from *edgeR*. Thirteen principal components (PC)s explaining more than 1% of the total variance were identified and their scores were tested against categorical variables of Batch, Sex and Line using separate 1-way ANOVA. An individual PC was considered significantly associated with a given categorical variable if the FDR corrected (a single variable across PCs) p-value was less than 0.05. The PC scores for each sample across all PCs, which explain at least 1% of the total variance, and the 1-way ANOVA test results for each biological variable of interest are included in Supplemental Table S6.

##### Note - Differential Genes and Co-Expression

The determination of DEGs requires low variance across samples, whereas high gene-gene correlations (used in WGCNA and DW) instead require variation across samples. Therefore, one expects gene sets generated from these methods to represent different expression regimes. However, given the high power (N = 193) of the present study, overlap of these gene sets is possible.

##### Differential Expression

Identification of DEGs was determined using the *limma + voom* version 3.50.0 pipeline in *edgeR* version 3.36.0. TMM normalized counts were variance corrected using the *voom* function and then fit to a model using line, sex and batch. Line, sex and batch were identified as the main sources of variance in the expression data using PCA analysis. A contrast was constructed to compare the high and low lines explicitly, collapsed on sex. A false discovery rate below 0.05, determined via the Benjamini-Hochberg (BH) procedure [99], was used to determine whether a given gene was DE. Statistics were calculated using the *eBayes* and *topTable* functions. In total, 2996 genes were identified as DE, with 1585 being down-regulated and 1411 being up-regulated in HP, relative to LP. DEG results are included in Supplemental Table S7.

##### Differential Variability

A DV gene is defined as having a significant (FDR < 0.05) difference in variance between groups. Each of the 10,716 genes included in WGCNA analysis was tested for DV using an F test (the *var.test* function in R’s base stats package), and FDR corrected using the BH method [99]. In total, 426 genes were identified as being DV (35 up and 391 down with respect to the HP line). Note that over 10x as many DV genes have a lower variability in the HP mice compared to the LP mice. DV results are included in Supplemental Table S8.

##### Differential Wiring

A DW gene is defined as having more differential edges than would be expected by chance, where a differential edge is a gene-gene correlation that changes by more than a threshold between the HP and LP lines. Briefly, the absolute difference in gene-gene correlation matrices is computed between the HP and LP lines and any change higher than a given threshold is tested for significance using the *r.*test function from the R psych package. Any edge with an FDR < 0.05 is defined as a differential edge. The threshold is arbitrary, but we find any choice above 0.5 that also limits the maximum number of differential edges per gene to below 100 works well in practice. Note that DW was developed by Dan Iancu [5, 6] to be a more conceptually (and computationally) approachable method of identifying gene networks and any methods to determine the cutoff threshold that involved clustering or network statistics would negate the purpose of the test. A cutoff threshold of 0.5 is generally well above the 99^th^ percentile and a choice that results in genes with more than 100 differential edges produces networks that are hard to interpret. The number of differential edges per gene is calculated using a row sum. A binomial test (R function *binom.test* in the stats package) is used to determine a p-value for each gene, with the number of trials being the total number of edges (number of genes squared), the probability of success being the number of differential edges / total number of edges, and an alternative hypothesis of “greater”. The results are FDR corrected using the BH method and any gene with an FDR < 0.05 was considered to be DW [99]. A total of 407 genes were identified as DW, with 284 having a net decrease in differential edge correlation, and 123 having a net increase in differential edge correlation (in HP compared to LP). DW results are included in Supplemental Table S9.

##### Weighted Gene Co-Expression Network Construction

Weighted gene co-expression networks were constructed following methods set forth by Langfelder and Horvath [20] and using the R package, *WGCNA* version 1.71. Following the WGCNA pipeline for unsigned networks established by Dan Iancu for the first HP vs LP selected lines, we drop genes with low connectivity [7, 8]. Unlike the previous pipeline, we additionally excluded non-protein-coding genes (n = 1,911) from further analysis in this work. Gene-level connectivity was determined by transforming gene-gene correlation matrices for the HP and LP mice into scale-free adjacency matrices. Briefly: (1) A soft thresholding power was determined by choosing a power that approximates a scale-free-network topology (R^2 > 0.8) but also maintains network median connectivity (10 for the HP network, 9 for the LP network); (2) The HP and LP networks were 95^th^ percentile quartile-normalized and then a network was constructed using the parallel max operation. This results in a network in which genes important to either of the HP or LP networks are maintained, hereafter referred to as a “max” network; (3) Gene-level network connectivity metrics were computed via a row sum on the max matrix; and (4) The lowest 10% of genes by connectivity (up to a maximum of 25% of all genes) were discarded prior to downstream analysis. In all cases within this work, the maximum 25% of genes (n = 3571) were discarded (on average corresponding to the lowest ∼1% of genes by connectivity), resulting in 10,716 genes remaining for analysis (all protein coding). The annotation for the dropped genes (non-protein coding and low connectivity) and the annotation for all genes included for further analysis are included in Supplemental Table S19. After removing low-connectivity genes, soft thresholding powers were re-calculated to verify that the optimal power did not change. Topological overlap matrices (TOMs) were then constructed from the adjacency matrices following the pipeline established by Langfelder and Horvath [20].

Once a TOM network is constructed, modules of genes are isolated using the dynamic tree cutting algorithm, *cutreeDynamic function,* on a hierarchical clustering built from the average distance of 1-TOM. We use the following arguments in the tree cutting function: *minClusterSize = 100, deepSplit = 4, respectSmallClusters = TRUE, pamStage = TRUE, pamRespectsDendro = FALSE.* We find these options, including the secondary partitioning around mediods (PAM) stage, to generate reproducible networks containing roughly 20 modules with a minimum gene membership of 100. Note, it is common to combine modules based on the correlation of their eigengenes, but we do not take that step. Modules are assigned an arbitrary color for identification. Note, when comparing network connectivity measures across the LP and HP networks, a soft power of 10 was used.

##### Hub Genes

Any gene in the top 20% of within-module connectivity is called a “hub gene”. These genes typically drive the ontology of a given module. Hub genes (n = 2138 HP, n = 2135 LP) were identified for all generated networks and tested for ontological enrichment, as discussed below in Section 4.4.6. We do not focus on a single hub gene per module, as in common (e.g. Salem et al. [93]), but that information is readily available in Supplemental Table S11 for the curious reader. Differences in the number of hub genes across networks result from rounding effects across modules with odd-number gene membership.

##### Unique Hub Genes

Of particular interest are genes that are hubs in the HP network, but not the LP network. These are called “unique HP hubs”. The opposite, genes that are hubs in the LP network but not the HP network, are called “unique LP hubs”. These correspond to genes that are driving connectivity and ontology in one network, but not another. Five hundred and sixty-two (562) unique hub genes were identified for the HP network, and 559 for the LP network.

Gene module membership (including hub and unique hub designations) is included for the HP, and LP networks in Supplemental Table S11. Summaries of each network’s modules, including enrichment (described below) p-values for cell-type, DE, DV, and DW genes, and unique hubs are included in Supplemental Table S12 for the HP, and LP networks at both the module and hub levels.

##### Preservation

There are many common definitions for network preservation, but for the sake of interpretability we have chosen to use Jaccard Similarity. The JS is the ratio of the number of terms in the intersection of two sets to the number of terms in the union of two sets, or in our case, the ratio of the number of items in common between two sets, and the total number of unique genes across both sets. For example, if you have two sets which are the same length, and have 66% of their items in common, the JS will be 0.4, and if they have 75% of their items in common the JS will be 0.5. We choose a middle value, JS > 0.45, as our cutoff for determining if two modules are preserved by comparing their hub genes.

#### 4.4.6 Over-Representation and Enrichment Analysis

##### Gene ontology

Enrichment of gene sets in GO Consortium terms was conducted using over representation analysis via the *enricher* function, which performs a hyper-geometric test, from the *clusterProfiler* v4.2.2 package. GO term annotation was assembled using the *AnnotationDbi* v1.56.2, *org.Mm.eg.db* v3.14.0, and *GO.db* v3.14.0 packages which were using the GO database from September 1st, 2021. The GO data used in this work is available at GitHub. Enrichment tests of hub and unique hub genes were carried out using the 10716 genes present in the WGCNA analysis as “universe genes”, all other enrichment tests used the 16,198 genes included in the DE analysis as “universe genes”, consistent with best practices [100]. A GO enrichment was considered significant if its FDR corrected p-value was < 0.05.

To reduce GO term redundancy, significant GO term results within a gene set were clustered based upon their JS, as discussed in Section 4.4.5. Clustering was done using tree cutting on hierarchical clustering based on a dissimilarity measure of (1 - JS) and a tree height of 0.3 (or a JS of 0.7, a much more stringent requirement than used in the network analysis). The GO term with the lowest FDR was kept as representative of the cluster.

For example, when looking at GO enrichment of the top 750 DEGs by absolute log fold change, three go terms GO:0030545 (signaling receptor regulator activity), GO:004818 (receptor ligand activity) and GO:0030546 (signaling receptor activator activity) had 21 genes in common (intersect) out of 23 unique genes (union) resulting in a JS of 0.91. These three GO terms were clustered. In general, a reduction of 1/3 of GO terms was common using this JS clustering.

##### Cell Type and other gene sets

Enrichment of identified genes (differential genes, module genes and hub genes) against other curated gene sets is carried out in the same methodology as above. Significant testing is done using a hyper-geometric test and FDR correction is carried out across modules. Cell type specific meta-markers were generated using markers identified during analysis of the snRNA-Seq data. First, meta-markers for inhibitory and excitatory types by taking the top 100 markers from the *coarse clustering* (see Section 4.4.3 for details) and eliminating genes if they were found in in more than two sets for maximum specificity. The same process was carried out for the *fine clustering* to identify highly specific meta-markers for inhibitory and excitatory sub-types, as well as for non-neuronal cell populations. Note that sub-type meta-marker enrichments were only ever considered in the event that the *overall* type (inhibitory or excitatory) was found to be significantly enriched. Meta-markers are available in Supplementary Table S20.

#### 4.4.7 Code and Data Availability

The data presented in this work are available in online repositories. Sequencing data (including fastq, raw and normalized counts) are available at the Gene Expression Omnibus with accession numbers GSE288772 (bulk) and GSE293636 (single nuclei). All code used to produce the manuscript, perform analysis or produce figures is available at the Portland Alcohol Research Center GitHub [https://github.com/parcbioinfo/HPLP_Preference_CeA], along with the raw counts, downstream analysis, and sample metadata needed to execute the scripts. Supplemental figures and tables results are also available at GitHub.

## Supporting information

Ontology tables for HP

Ontologies for LP

## 5 Acknowledgement

This work was supported by the National Institute on Alcohol Abuse and Alcoholism (P60 AA010760, T32 AA007468, L70 AA031860, U01 AA013519, R01 AA026278, R01 AA027552), the US Department of Veterans Affairs (I01BX004699, I01BX006570, IK6BX006342) and a generous gift from the John R. Andrews family. The contents do not represent the views of the U.S. Department of Veterans Affairs or the United Stated Government. The research reported in this publication used computational infrastructure supported by the Office of Research Infrastructure Programs, Office of the Director, of the National Institutes of Health under Award Number S10OD034224. The content is solely the responsibility of the authors and does not necessarily represent the official views of the National Institutes of Health.

## Notes

https://www.ncbi.nlm.nih.gov/geo/query/acc.cgi?acc=GSE288772

https://www.ncbi.nlm.nih.gov/geo/query/acc.cgi?acc=GSE293636

https://github.com/parcbioinfo/HPLP_Preference_CeA

### Competing Interest Statement

The authors have declared no competing interest.

