## Supplemental Figure Captions for "Genomic and Behavioral Signatures of Selection for Ethanol Preference from the Heterogeneous Stock Collaborative Cross Mice – The Central Nucleus of the Amygdala"

**JQA_HPLP_CeA_Supplemental Info_submitted.docx**

This contains additional behavioral results, as well as all supplemental figures which are referenced within the main text.

**JQA_HPLP_CeA_Supplemental Tables_submitted.xlsx**

This contains all supplemental tables which are referenced in the text, and a table of contents.

**JQA_HPLP_CeA_Supplemental Ontology Tables_HP.xlsx**

**JQA_HPLP_CeA_Supplemental Ontology Tables_LP.xlsx**

These files contain enriched Gene-ontologies for each of the modules hug genes which were identified from weighted gene co-expression network analysis on either the high-preferring (HP) or low-preferring (LP) mice.
