## Supplemental Info for "Genomic and Behavioral Signatures of Selection for Ethanol Preference from the Heterogeneous Stock Collaborative Cross Mice – The Central Nucleus of the Amygdala"

### 1 Two-bottle choice drinking of saccharin, quinine and sucrose

#### 1.1 Rationale and Methods

Because taste and differential intake of natural rewards could be factors that impact ethanol drinking, we examined the intake of flavored solutions (saccharin, quinine) and a natural caloric reward (sucrose) in the HP and LP mice. Testing for the intake of bitter- and sweet-flavored solutions is common practice in rodents bred for differences in ethanol intake. In prior studies, differences in quinine intake rarely have been found. A relationship of higher intake of sweet solutions in high compared to low ethanol consuming lines has more often been found (Agabio et al., 2000; Grahame et al., 1999; Phillips et al., 2005; Sinclair et al., 1992; Stewart et al., 1994). Therefore, we predicted no difference in quinine consumption between the HP and LP lines and greater intake of saccharin and sucrose in the HP compared to LP line. Data were collected for each flavored solution in a single group of mice with the order in which the flavors were offered counterbalanced. Each solution was offered vs. a water choice for 8 days for 24 h per day with a lower concentration offered for the first 4 days followed by a higher concentration. These methods are consistent with those used previously (Crabbe et al., 1996; 2011; Reed et al., 2021) and reduce animal usage by testing all types of solutions in one set of mice. Methods for measuring intake via calibrated drinking tubes were the same as described in the main manuscript for measuring ethanol intake.

Data for each type of solution were analyzed by ANOVA, with concentration as a repeated measure and selected line and sex as independent variables. As for ethanol drinking, the relative positions of the water and flavored solution tubes were switched every two days to control for side preferences. The dependent measures for each solution were preference ratio (ml of flavor consumed/ total fluid consumed in ml), mg/kg intake, and ml/kg total volume consumed, based on the average of days 2 and 4 and of days 6 and 8, which were the second days after a tube position switch for each concentration.

#### 1.2 Results

##### 1.2.1 Sweet Solutions

**Saccharin:** Repeated measures ANOVA identified a significant line by concentration interaction for both preference (F(1,44)=8.7, p<0.01) and intake (F(1,44)=7.9, p<0.01). HP mice exhibited greater preference than LP mice for the low saccharin concentration (Figure S2A) and consumed more saccharin at both saccharin concentrations, compared to LP mice (Figure S2B). Preference for the two saccharin concentrations was comparable for HP mice, but greater for the higher concentration in LP mice. Consumption of saccharin was greater when offered at the higher concentration for both mouse lines. For total volume consumed (Figure S2C), there was also a significant line by concentration interaction (F(1,44)=4.2, p<0.05). HP mice consumed more total volume than LP mice at both concentrations of saccharin, and both lines increased total volume consumed at the higher saccharin concentration. There were no significant effects of sex for preference, consumption or total volume.

**Sucrose:** On some days, mice consumed all of the sucrose solution available. In these cases, a consumption volume of 25 ml was assigned. Repeated measures ANOVA identified no effects of line for sucrose preference (Figure S3A), but there was a significant increase in preference at the higher concentration (F(1,44)=8.2, p<0.01). For sucrose consumption, there was a significant line by concentration interaction (F(1,44)=6.0, p<0.05). HP mice consumed more sucrose than LP mice at both sucrose concentrations (ps<0.01) and both selected lines consumed more sucrose when it was offered at the higher concentration (ps<0.001) (Figure S3B). There was a significant main effect of sex for both preference (F(1,44)=7.3, p<0.01) and consumption (F(1,44)=5.8, p<0.05). Females exhibited higher preference for sucrose than males (mean ± SEM = 0.95 ± 0.01 vs. 0.91 ± 0.01 for the preference ratio for females and males, respectively), as well as greater intake (mean ± SEM = 15.3 ± 11.7 g/kg vs. 12.8 ± 1.4 g/kg for females and males, respectively), regardless of concentration. For total volume consumed (Figure S3C), there were significant main effects of line (F(1,44)=21.3, p<0.001) and concentration (F(1,44)=119.5, p<0.001). HP mice consumed more total volume than LP mice and total volume increased across concentrations. There were no significant effects of sex for total volume.

**Conclusions:** The greater saccharin and sucrose intake of mice that consume more ethanol is consistent with several prior studies in selected rodent lines (Grahame et al., 1999; Phillips et al., 2005; Sinclair et al., 1992; Stewart et al., 1994), as well as in other mouse populations such as panels of standard inbred strains and among individuals of a heterogeneous stock (Belknap et al., 1993; Overstreet et al., 1993; Yoneyama et al., 2008). These data suggest that some common genes influence ethanol and sweet solution intake.

##### 1.2.2 Bitter Solution

**Quinine:** Repeated measures ANOVA identified no significant line differences for quinine preference (Figure S4A), consumption (Figure S4B) or total volume (Figure S4C). There were significant main effects of concentration for preference (F(1,44)=13.4, p<0.001), amount of quinine consumed (F(1,44)=18.7, p<0.001), and total volume consumed (F(1,44)=8.3, p<0.01). Mice exhibited lower preference for the higher quinine concentration but had higher intake and total volume at the higher quinine concentration. There were no significant effects of sex for quinine preference or intake but there was a main effect of sex for total volume (F(1,44)=4.8, p<0.5), with females consuming more total volume per kg body weight than males (mean ± SEM = 252.8 ±13.5 ml/kg and 220.8 ± 9.0 ml/kg for females and males, respectively).

**Conclusions:** Intake and preference for a bitter-tasting solution do not differ between the selectively bred HP and LP lines, indicating that the bitter taste quality of ethanol is not a likely factor in their differential ethanol intake and preference.

### 2 Ethanol-induced locomotor stimulation

#### 2.1 Rational and Methods

Ethanol has time- and dose-dependent effects on locomotor activity. Locomotor stimulant effects in mice occurring at early periods after exposure or in response to lower ethanol doses model ethanol-induced behavioral stimulation in humans, whereas reduced locomotor activity after higher doses of ethanol models behavioral depression. Some, though not all, human data support an association of high stimulation and low behavioral depression in response to ethanol with higher risk for ethanol use (Chavarria et al., 2021; King et al., 2016; Lippard et al., 2023; Plawecki et al., 2022). We examined this in the HP and LP lines, hypothesizing that the HP mice would exhibit greater locomotor stimulation and less locomotor depression, compared to the LP mice.

Mice were tested on three consecutive days as described previously (Gubner et al., 2013; Kamens et al., 2006; Palmer et al, 2002) in 16 automated SuperFlex activity monitors (OmniTech Electronics; Columbus, OH) that utilize photocell beam interruptions to measure distance traveled. Each monitor was equipment with 16 infrared photocell beams located 1.6 cm above the floor of a 40 cm x 40 cm x 30 cm clear acrylic chamber. Environmental control chamber enclosures surrounded each monitor (AccuScan Instruments, Inc., Columbus, OH), and were each illuminated by a 3.3 W incandescent light bulb and equipped with a fan that provided ventilation and background noise.

On each test day, mice were transported to the procedure room and allowed to acclimate for 45 minutes. They were then weighed and placed into holding cages while syringes for injections were filled. Saline or ethanol was then administered (IP) and each mouse was immediately placed into the activity monitor for 15 minutes. Data were collected in 5-min time periods. On days 1 and 2, all mice received a saline injection, while on day 3 mice received an injection of saline or ethanol (1.0, 1.5, 2.0 or 2.5 g/kg; 20% v/v ethanol solution). Day 1 of testing familiarized the animals with the testing procedure and environment, and day 2 of testing provided a measure of baseline activity. To account for baseline differences in the calculation of the locomotor effects of ethanol, day 2 baseline distances for individual mice were subtracted from each individual’s saline or ethanol day 3 distance. This difference score was used as the dependent variable for each 5-min period, consistent with our previous work (Gubner et al., 2013; Kamens et al., 2006; Palmer et al., 2002; Phillips et al., 1995). Immediately after testing on day 3, a 20 µl blood sample was obtained from the retro-orbital sinus to determine blood ethanol concentration (BEC; mg/ml) using gas chromatography (Finn et al., 2007; Rustay and Crabbe, 2004).

#### 2.2 Results

Data were examined by repeated measures ANOVA for changes across 5-min time periods, with line and sex as grouping factors. There were significant main effects of line (F[1,171]=4.1, p<0.05), dose (F[4,171]=8.4, p<0.0001) and time (F[2,342]=3.5, p<0.05). There was no main effect of sex and sex did not interact with line or dose. To examine time-dependent effects, we examined the data for each 5-min period. There was a significant line x dose interaction for the first 5 minutes (F[4,171]=3.0, p<0.05). The LP mice exhibited greater stimulation than HP mice after treatment with the 2 g/kg ethanol dose (Figure S5A). Simple main effect analyses indicated that both lines exhibited dose-dependent effects (p<0.05 for HP mice and p<0.001 for LP mice). Follow-up mean comparisons (Tukey HSD) identified significantly greater stimulation by the 2.5 g/kg ethanol dose, compared to saline in HP mice and by the 1.5, 2.0 and 2.5 g/kg ethanol doses, compared to saline in LP mice. For the second 5-min period, there were significant main effects of line (F[1,171]=4.6, p<0.05) and dose (F[4,171]=6.8, p<0.0001), but no line by dose interaction. Overall, LP mice were more stimulated than HP mice (Figure S5B). For the third 5-min period, there was a significant dose effect (F[4,171]=5.6, p<0.001), but no effect of line. Neither line exhibited locomotor depression to this range of ethanol doses during the 15-min test period (Figure S5C). Analysis of BEC in samples obtained immediately after ethanol-induced locomotor activity testing supported dose-dependent BECs (F[3,144]=141.9, p<.0001) but revealed no significant line difference (Figure S5 inset). There were no sex effects for BEC.

**Conclusions:** These data indicate that sensitivity to the locomotor stimulant effects of ethanol in this set of selected lines corresponds with lower, rather than higher, ethanol intake. Differences in locomotor response to ethanol were not related to differences in BEC. It’s possible that greater sensitivity to ethanol’s stimulant effects in LP mice plays a role in their lower voluntary ethanol consumption.

### 3 Ethanol-induced hypothermia

#### 3.1 Rational and Methods

Ethanol lowers core body temperature, producing hypothermia. Mixed results have been obtained for studies examining a potential genetic relationship between ethanol drinking and sensitivity to the hypothermic effects of ethanol (Crabbe et al., 2012; Cunningham et al., 1991; Gehle and Erwin, 1998; Stewart et al., 1992; Tampier and Mardones, 1999). We examined this in the HP and LP mice after an acute injection of 2 or 4 g/kg ethanol (IP; 20% v/v). Our standard testing procedures were used (Harkness et al., 2015; Palmer & Phillips, 2002; Shen et al 1996). Mice were transported to the experiment room, weighed, and placed into cubicles within customized perforated acrylic plastic chambers designed to prevent huddling, so that accurate measures of body temperature can be obtained. Mice were left to acclimate for one hour and then body temperatures were obtained at time (T) 0, 30, 60, 90, 120, 180, 240 and 300 minutes, using a glycerol-lubricated rectal probe (1.2 mm ball x 2 cm length; Sensortek Thermalert Th-8, Bailey Instruments, Inc., Saddle Brook, NJ). The T0 temperature was taken just prior to IP injection with saline or ethanol and mice were replaced back into their cubicles between temperature assessments.

#### 3.2 Results

In a repeated measures ANOVA with time as the repeated measure and line, sex and dose as factors, no significant main or interaction effects involving line were found. There were significant main effects of sex (F[1,107]=4.4, p<0.05), dose (F[2,107]=249.5, p<0.0001), and time (F[7,749]=110.6, p<0.0001) and a significant dose by time interaction (F[14,749]=64.1, p<0.0001). Overall, males were colder than females by about 0.2°C, but this difference was not dependent on ethanol dose or time. Data are graphed in 3 panels by HP (Fig. S6A), LP (Fig. S6B) and combined (Fig. S6C), since there were no significant line effects. Because there were no line differences, simple effects analysis and mean comparisons were carried out only for the combined data (Fig. S5C). There were significant effects of dose at all timepoints except T0 (ps<0.001). Mean body temperature of mice treated with 2 g/kg ethanol was significantly lower than that of the saline-treated controls at T30-120. Mean body temperature of mice treated with 4 g/kg ethanol was significantly lower than that of both the saline- and 2 g/kg-treated mice at T30-300. The simple effect of time was significant for each dose group (ps≤0.001). Data were next examined for temperature changes from one time measurement to the next. In the saline group, temperature dropped significantly only from T120 to T180; a drop of 0.3 °C. For the 2 g/kg ethanol group, maximum hypothermia occurred at T30; a drop of 1.6°C from T0. For the 4 g/kg ethanol group, the hypothermic response continued through T120 and then began to recover. The maximum drop in mean body temperature for the 4 g/kg ethanol dose was 4.9°C at T120 compared to T0.

**Conclusions:** These data indicate that selective breeding for differential ethanol intake did not result in differential sensitivity to the hypothermic effects of ethanol and thus, that common genes do not impact these two ethanol traits in this pair of selected lines.

### 5 Supplemental Figures

#### 5.1 S1 – Intake and Preference Correlations

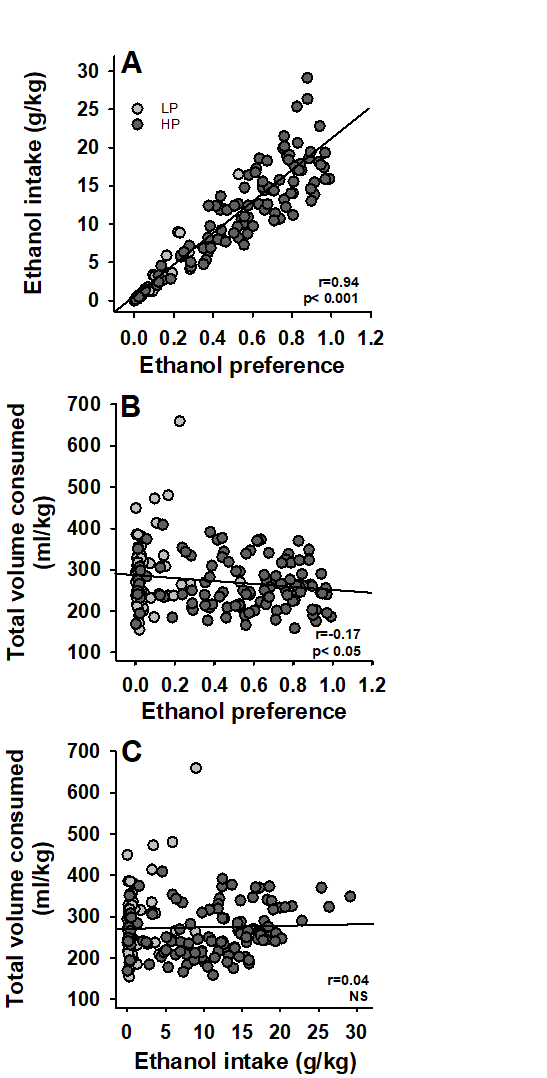

**Figure S1 - Phenotypic correlations:** Shown are correlations between ethanol preference, ethanol intake and total volume consumed for S4 LP and HP offspring. N=174 (63 LP and 111 HP mice).

#### 5.2 S2 - Saccharin

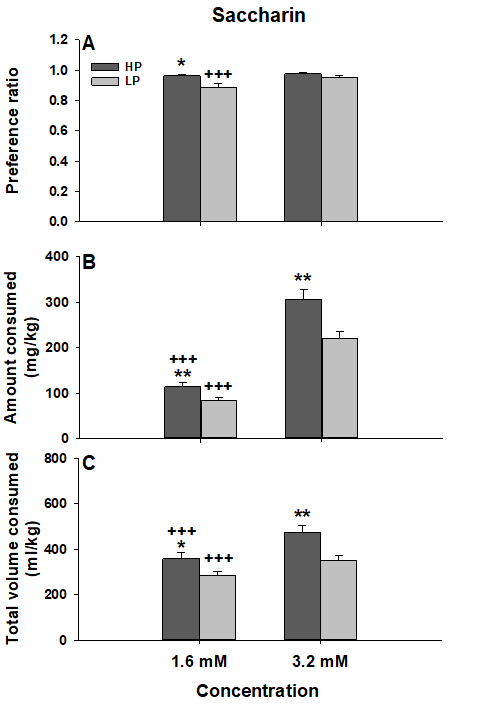

**Figure S2 - HP mice exhibit greater saccharin preference and intake, compared to LP mice:** Shown are means ± SEM for S5 generation HP and LP mice. (A) Preference ratio (ml from saccharin tube / total ml); (B) Saccharin intake (mg/kg); (C) Total volume consumed (ml/kg) from the water and saccharin tubes. N=24 HP (12 female and 12 male) and 24 LP (12 female and 12 male) mice; mean age (± SEM) at the beginning of testing was 97 ± 0.7 days (range = 91-101 days). *p<0.05, **p<0.01 for the difference between HP and LP mice; +++p<0.001 for the effect of concentration.

#### 5.3 S3 - Sucrose

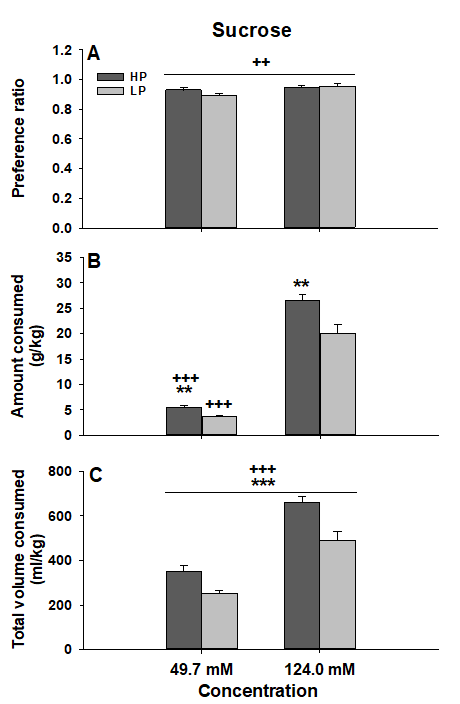

**Figure S3 - HP mice exhibit greater sucrose intake, but not preference, compared to LP mice:** Shown are means ± SEM for S5 generation HP and LP mice. (A) Preference ratio (ml from sucrose tube / total ml); (B) Sucrose intake (g/kg); (C) Total volume consumed (ml/kg) from the water and sucrose tubes. Data were collected for the same mice from which saccharin drinking data were collected. **p<0.01, ***p<0.001 for the difference between HP and LP mice; ++p<0.01, +++p<0.001 for the effect of concentration.

#### 5.4 S4 - Quinine

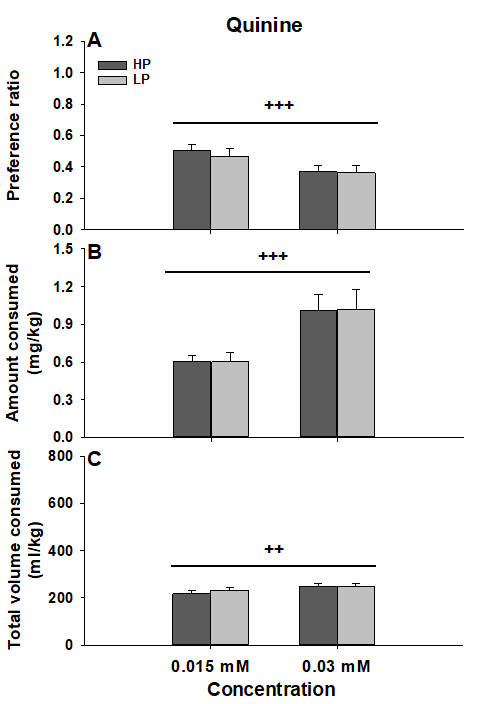

**Figure S4 -** **HP and LP mice exhibit similar quinine preference and intake:** Shown are means ± SEM for S5 generation HP and LP mice. (A) Preference ratio (ml from quinine tube / total ml); (B) Quinine intake (mg/kg); (C) Total volume consumed (ml/kg) from the water and quinine tubes. Data were collected for the same mice from which saccharin and sucrose drinking data were collected. ++p<0.01, +++p<0.001 for the effect of concentration.

#### 5.5 S5 - Locomotor

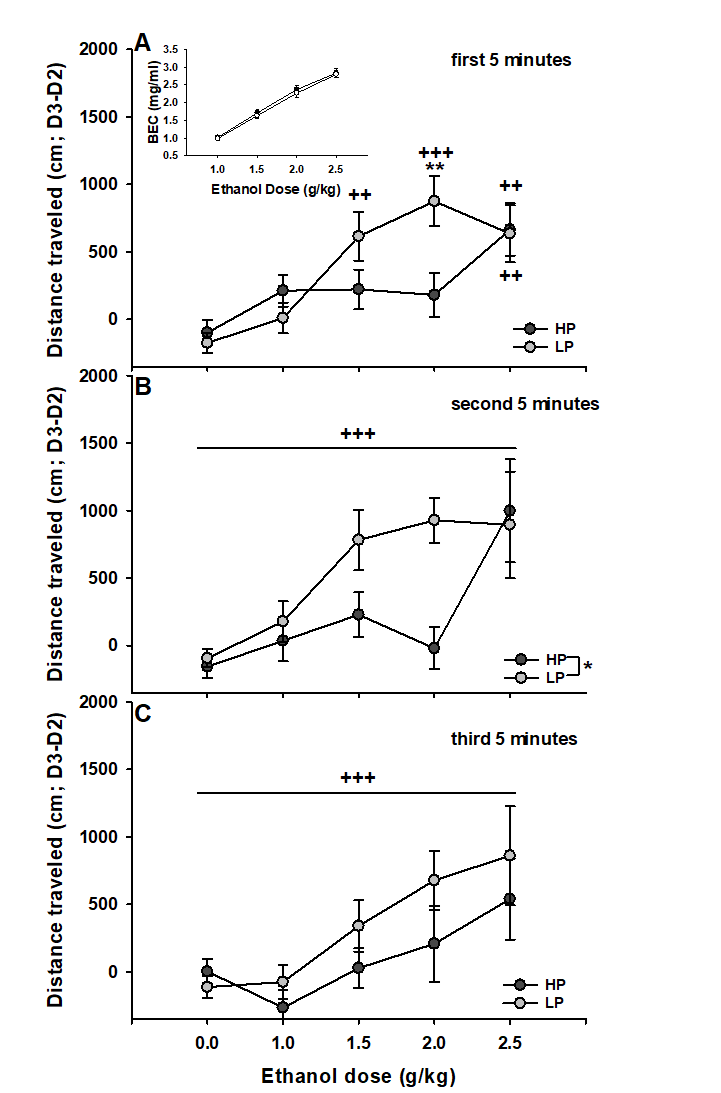

**Figure S5 - LP mice are more sensitive to the locomotor stimulant effects of ethanol than HP mice:** Shown are means ± SEM distance traveled after saline (0.0) or ethanol (1.0 - 2.5 g/kg) treatment corrected for baseline distance (D3-D2) for three consecutive 5-minute test periods (A-C). The inset shows blood ethanol concentrations (BEC; mg/ml) taken from the retro-orbital sinus immediately after the total 15-min test. N=191 S5 HP and LP mice (6-10/sex/line/dose); mean age (± SEM) at the beginning of testing was 88 ± 5 days (range = 76-95 days). *p<0.05, **p<0.01 for the effect of line; ++p<0.01, +++p<0.001 for the main effect of dose or of the indicated dose compared to 0.

#### 5.6 S6 - Hypothermia

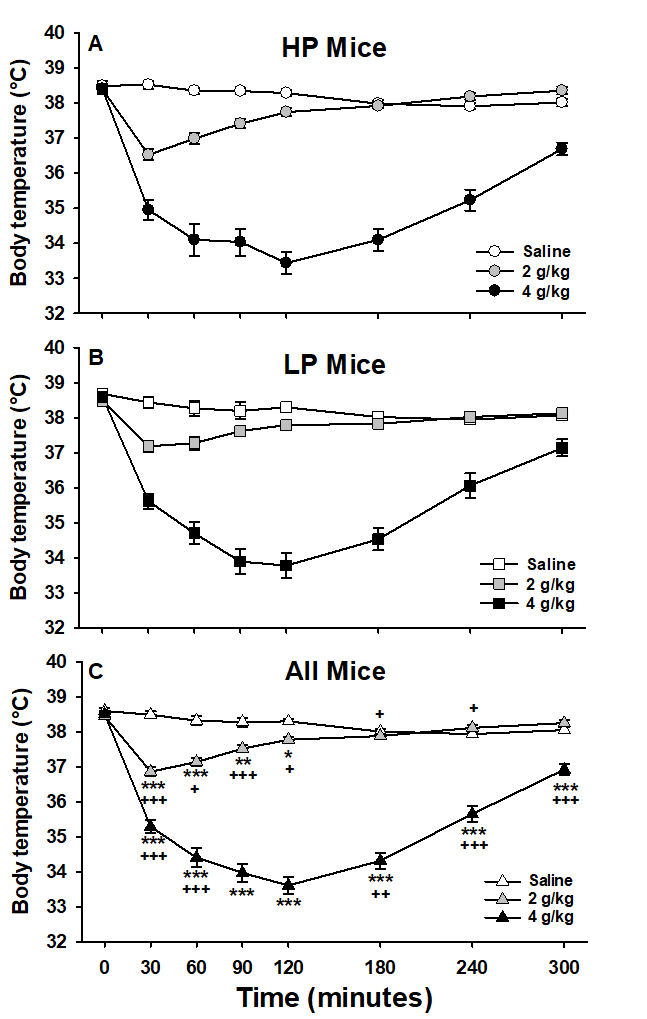

**Figure S6 -** **HP and LP mice do not differ in thermal response to ethanol:** Shown are means ± SEM body temperature after saline or ethanol treatment (2 or 4 g/kg) in S5 HP (A) and LP (B) mice, and all mice combined (C). Hypothermic response to ethanol in S5 HP and LP mice. Mean comparisons were conducted only for the combined data because there were no line differences. N=120 (10/sex/line/dose), 62-101 days of age. *p<0.05, **p<0.01, ***p<0.001 for the effect of dose compared to saline; +p<0.05, ++p<0.01, +++p<0.001 for change in body temperature at a given time, compared to the previous time within a given dose.

#### 5.7 S7 – Cell type Composition

A)

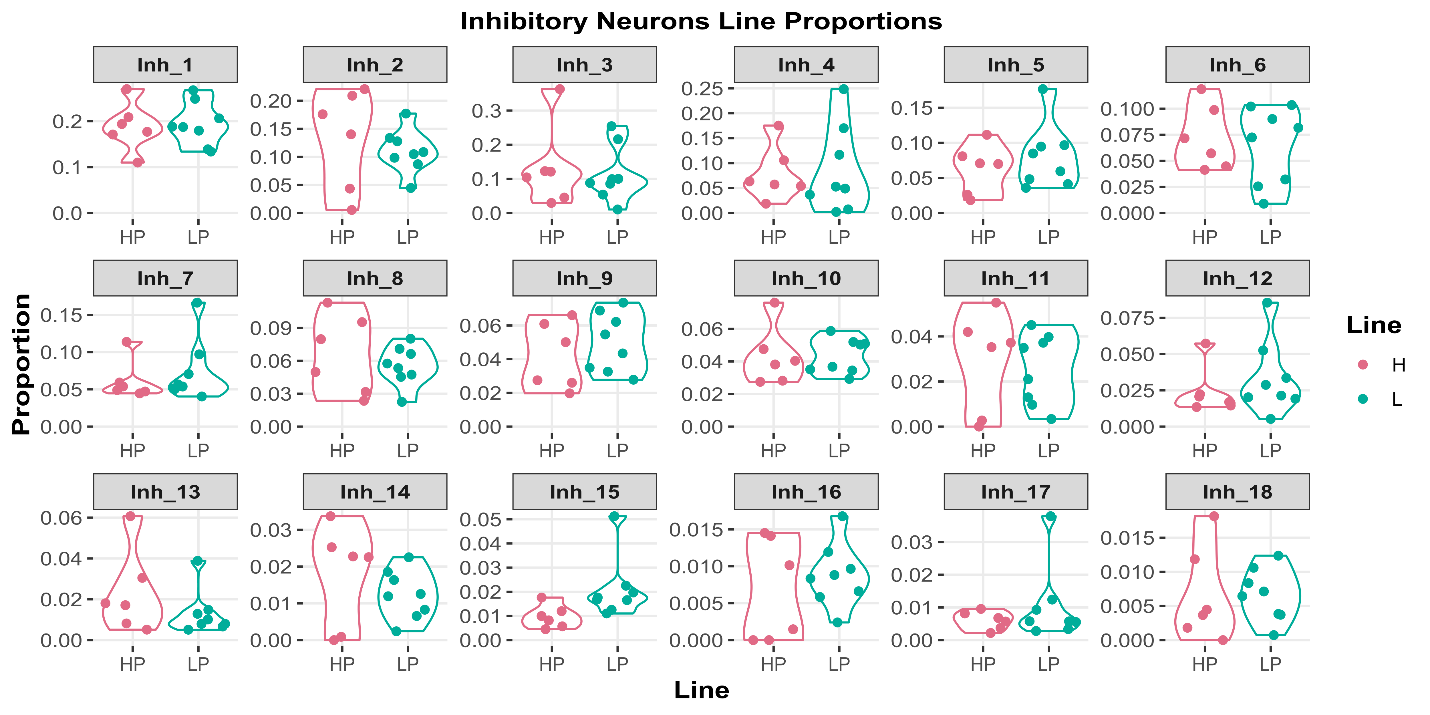

B)

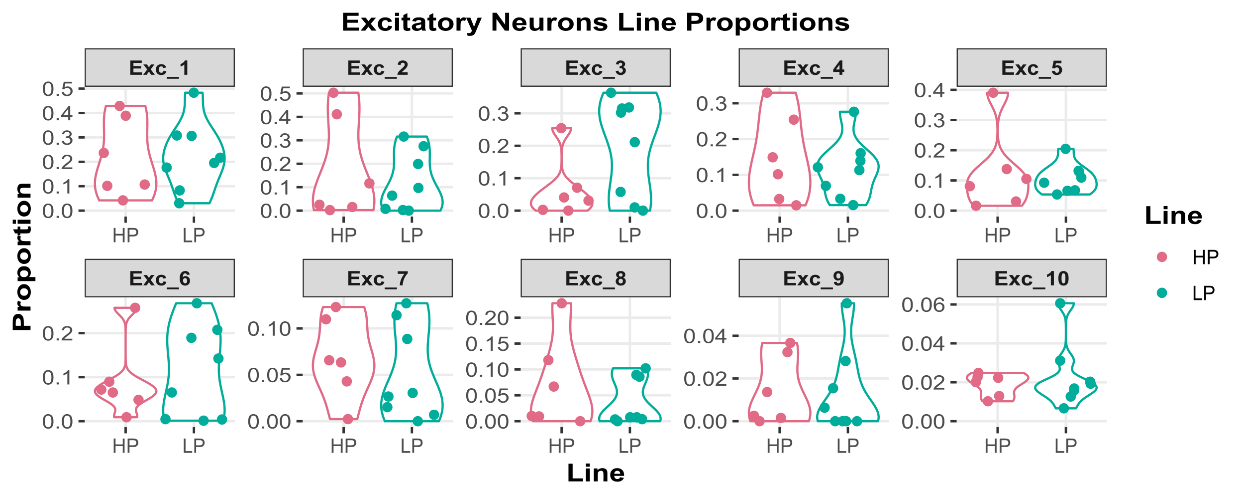

C)

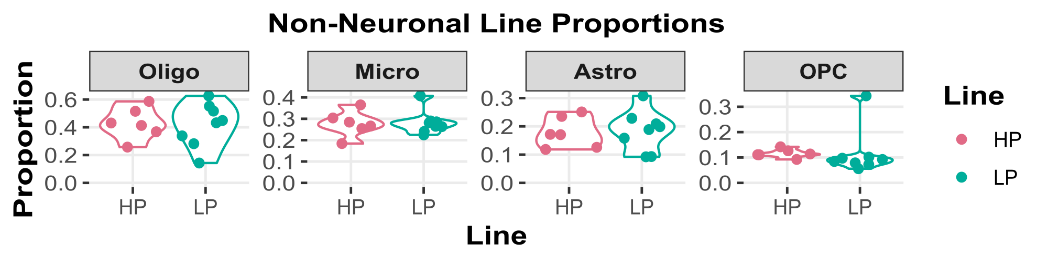

**Figure S7 -** Proportional composition of the HP and LP lines for each of the identified (A) inhibitory neuronal, (B) excitatory neuronal and (C) non-neuronal sub-types. Each individual animal included in analysis (N=14) is shown. No significant differences were observed for any sub-type.

#### 5.8 S8 – Excitatory Neuronal DEGs

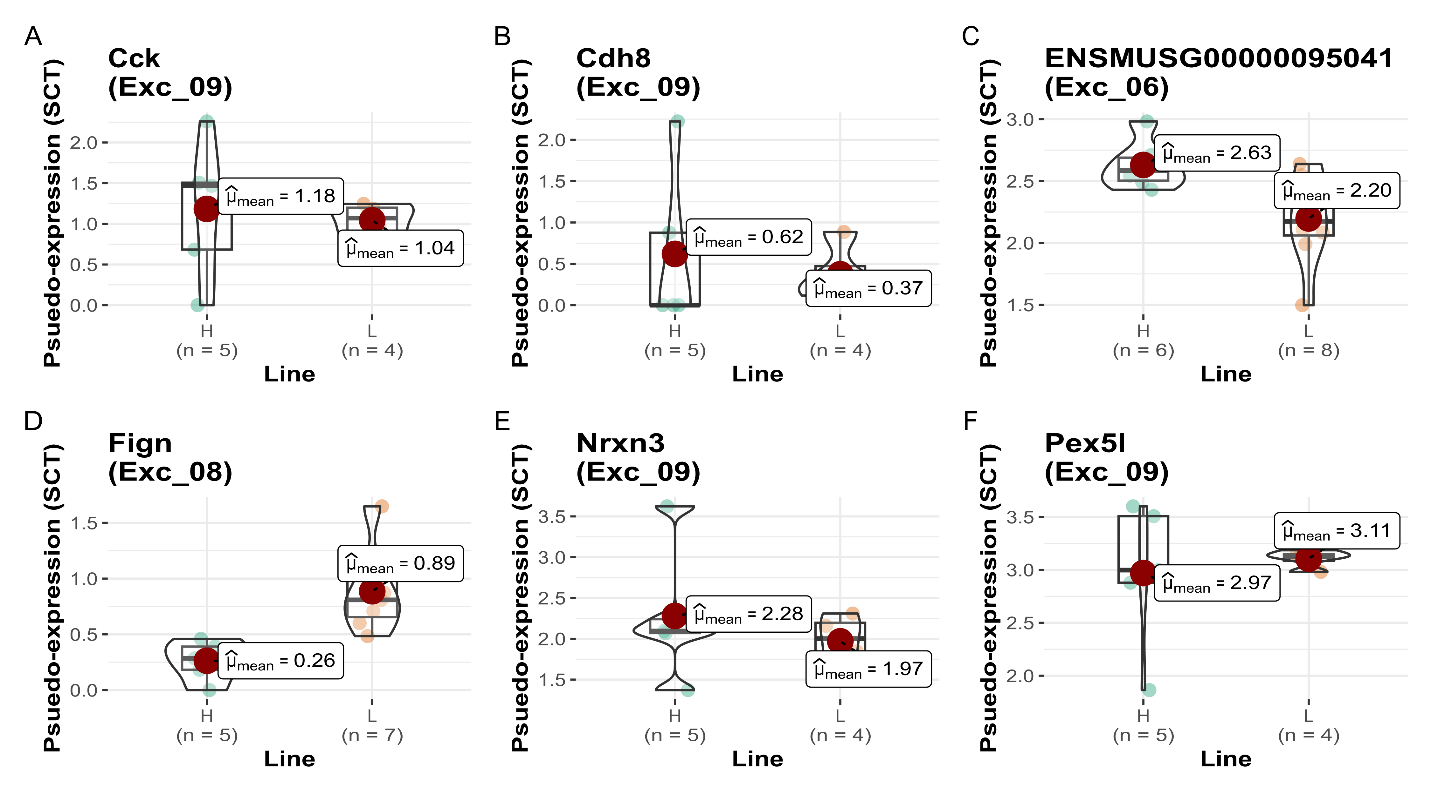

**Figure S8 -** Psuedo-bulk normalized expression levels for all genes found to be significantly differentially expressed in the excitatory neuronal sub-types. All four genes from Exc_9 [*Cck* (A), *Cdh8* (B), *Nrxn3* (E), and *Pex5l* (F)] appear to be driven by outliers. *Fign*, in Exc_08, is downregulated in the HP line, while the unannotated pseudogene *ENSMUSG00000095041* from Exc_06 is upexpressed in the HP line.

#### 5.9 S9 – bulkRNA-Seq PCA

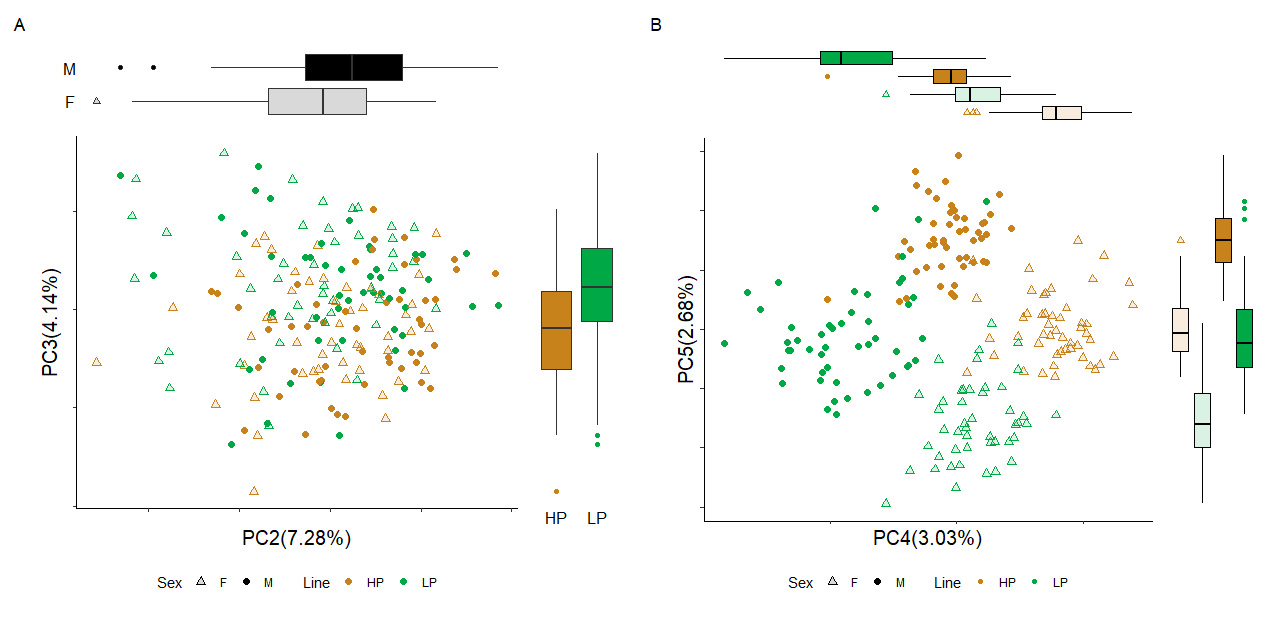

**Figure S10** - Principal Component (PC) Analysis – PC scores for each of the 193 animals included in the analysis using *limma + voom* normalized counts for the 16198 genes included in analysis. Green corresponds to low-preference, and orange corresponds to high-preference mice. Solid colors (circles) correspond to males, and light colors (triangles) corresponds to females. (A) PC scores for components 2 and 3 that together explain 11% of the variance present in the expression data. PC2 is significantly associated with sex (FDR < 4e-3), and PC3 is significantly associated with line (FDR < 4e-6). (B) Scores for PCs 4 and 5 that together explain about 5.5% of total variance present in the expression data and are both (FDR < 3e-40) significantly associated with Sex x Line. Each of PCs 4 and 5 group across line and sex, with PC5 separating male HP mice from female LP mice, while PC4 separate male LP mice from female HP mice. The variance explained by sex and line are additive, with no interaction observed. Boxplots are included for each PC, with counts grouped by the categorical variable most significantly associated with the PC. Significance was determined via FDR corrected one-way ANOVA.

#### 5.10 S10 – snRNA-seq dropped samples composition

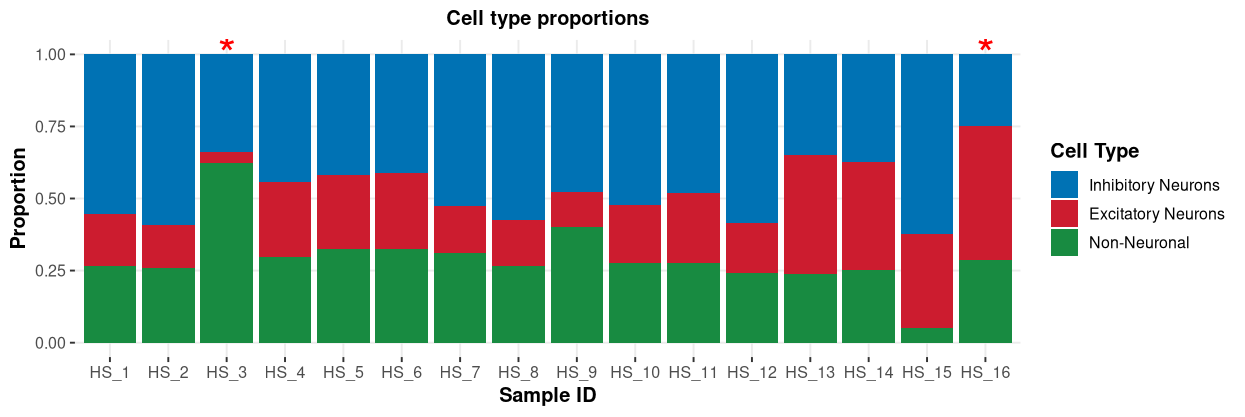

**Figure S10 – Broad cell-type composition of snRNA-seq samples:** After cell-type was determined using identified cell markers and proportions of Inhibitory (blue) and excitatory (red) neurons, as well as non-neuronal cells (green) were determined for all 16 samples which underwent snRNA sequencing. Two samples were identified as abnormal and removed from additional analysis (red star): HS_3 had ~2x the proportion of non-neuronal cells, and HS_16 had a ~2x lower ratio of inhibitory to excitatory neurons. An additional sample, HS_15, had a low count of non-neuronal cells, but a nominal ratio of inhibitory to excitatory neurons so was not excluded.

#### 5.11 S11 – bulkRNA-Seq zero gene distributions

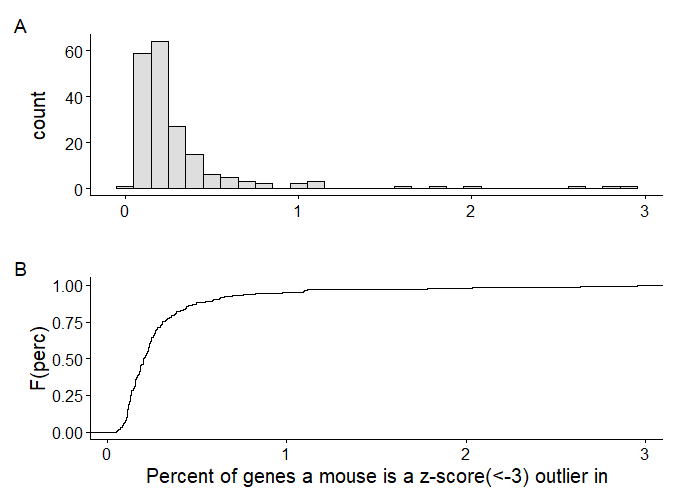

**Figure S11 -** Distribution of mice based on the percentage of genes they are low-count (TMM normalized) outliers in, based on having a z score less than -3. A) A histogram showing the number of mice with a given percent of genes they are low-count outliers in, B) an empirical-cdf showing the proportion of total mice which have less than a given percent of genes they are low-count outleirs in.

#### 5.12 S12 – bulkRNA-Seq zero gene distributions

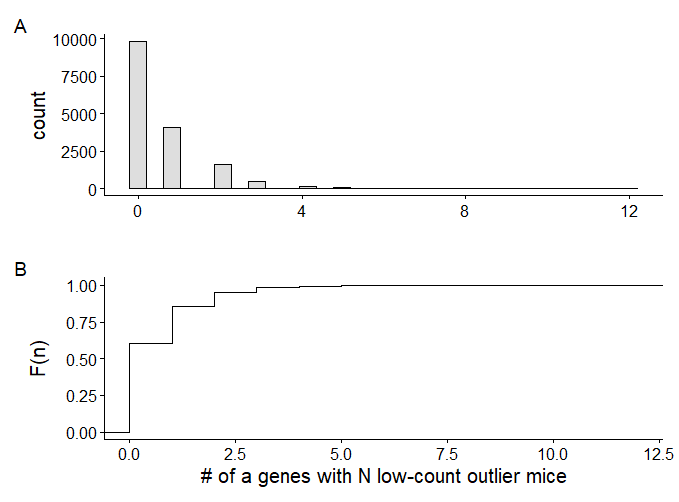

**Figure S12 -** Distribution of genes based on the number of mice which are low-count (TMM normalized) outliers, based on having a z score less than -3. A) A histogram showing the number of genes with a given number of low-count outlier mice, B) an empirical-cdf showing the proportion of total genes which have less than a given number of low-count outlier mice.
